# A GPCR-G protein-β-arrestin megacomplex induced by an allosteric modulator

**DOI:** 10.1101/2025.03.24.645131

**Authors:** Guodong He, Qinxin Sun, Xinyu Xu, Fang Kong, Shuhao Zhang, Kexin Ye, Xiaoou Sun, Xin Chen, Chuangye Yan, Xiangyu Liu

## Abstract

GPCRs signal through both G protein pathways and β-arrestin pathways. Previous work suggests that β-arrestins bind to GPCRs through different modes, including core engagement and tail engagement. Core engagement competes with G proteins and terminates G protein signaling, while tail engagement can coexist with G proteins, mediating sustained intracellular activation of the receptor – a process dependent on the high affinity between β-arrestin and the phosphorylated C-terminus of the receptor. In this study, we determined the structure of a GPCR - G protein - β-arrestin-1 complex stabilized by an allosteric modulator. The compound, atazanavir, acts like molecular glue to anchor β-arrestin-1 to the receptor’s TM6 and TM7 regions. This ‘pendulum’ binding mode is structurally compatible with simultaneous G protein binding. We further demonstrate that the atazanavir-mediated β-arrestin recruitment does not require the receptor’s C-terminal region. This work illustrates a novel paradigm of GPCR-G protein-β-arrestin1 megacomplex assembly and opens up new avenues for modulating GPCR function.

## Introduction

G protein-coupled receptors (GPCRs) mediate the perception of extracellular signals and the regulation of cellular activity. Upon binding to an agonist, GPCRs initiate G protein signaling, whose magnitude and duration are tightly regulated by β-arrestin (βarr)-mediated desensitization processes. This process involves receptor phosphorylation by the GPCR kinases (GRKs) and the recruitment of βarrs to the phosphorylated receptor. Additionally, βarrs also function as adaptors to mediate distinct βarr-associated signaling pathways^1^. For some GPCRs, biased ligands that preferentially activate G protein signaling or βarr signaling may exhibit better pharmacological properties^2^.

Due to their significant roles in regulating physiological responses and their value as drug targets, structural biology studies of GPCRs have attracted widespread attention^3,4^. The structure of the β_2_-adrenergic receptor (β_2_AR)-Gs complex, reported in 2011, was the first to capture how a GPCR transmits extracellular signals to intracellular G proteins through conformational changes^5^. The rhodopsin-βarr complex structure, reported in 2015, was the first to reveal how βarr recognizes GPCRs^6^. In the structures, G protein and βarr bind to the same intracellular pocket on the GPCR, indicating a competitive relationship between them. These structures nicely explain how βarrs compete with G proteins and terminate G protein signaling^1,5–7^.

However, the interaction between βarrs and GPCRs is more complex than what was revealed by the rhodopsin-βarr structure^6^. Previous studies have reported that some GPCRs exhibit sustained intracellular signaling^8–10^. These GPCRs are characterized by a ‘Class B’ mode of βarr binding, where the GPCR maintains a continuous, high-affinity interaction with βarr^11,12^. This phenomenon has led to the hypothesis that βarrs can coexist with G proteins, suggesting the possible existence of GPCR-G protein-βarr megacomplexes^8,13^. A breakthrough in this area came from the Lefkowitz laboratory, which first reported the structure of a megacomplex where a β_2_V_2_R chimeric receptor simultaneously binds to Gs protein and βarr^13^. The β_2_V_2_R chimeric receptor was constructed by fusing the V_2_R C-terminus, which binds strongly to βarr, to the β_2_AR. Though this strategy, researchers captured a βarr binding mode that does not compete with G proteins^13^, termed the ‘tail engagement’ mode. On the contrary, the βarr binding mode captured by the rhodopsin-βarr complex was named ‘core engagement’, characterized by βarr binding to the core of the intracellular ends of the transmembrane helices^6^. Recently, Chen et al. also reported a different ‘tail engagement’ binding mode for the glucagon receptor with βarr^14^. Structural comparisons revealed that in this binding mode, βarr does not compete with the alpha subunit of G proteins, suggesting a binding state that may coexist with G proteins. Notably, in this work, researchers also employed the strategy of fusing the V_2_R C-terminus to enhance the affinity between the receptor and βarr^14^.

These two pioneering studies demonstrated the possibility of GPCRs simultaneously binding to both G proteins and βarrs^13,14^. In both studies, a C-terminus with high affinity for βarr appears to be essential for the formation of such a megacomplex, as both structures utilized the C-terminus of V_2_R, known for its high affinity for βarr^13–17^. This also suggests that the formation of GPCR - G protein - βarr megacomplexes is likely an intrinsic property of GPCRs with ‘Class B’ βarr binding modes^18^. In our companion paper, we reported the discovery of atazanavir as a pan-positive allosteric modulator (’pan-PAM’) that can activate a range of GPCRs. More intriguingly, atazanavir mediates sustained intracellular activation of ‘Class A’ GPCRs, which have low affinity for βarr^19^. Further biochemical experiments showed that atazanavir mediates the formation of a GPCR-G protein-βarr complex. To gain deeper insights into the molecular mechanism by which atazanavir achieves its activity, we determined the high-resolution structure of an atazanavir-mediated GPCR-G protein - βarr megacomplex. As detailed below, this structural study did not employ the strategy of fusing the V_2_R C-terminus. Further studies even demonstrated that atazanavir-mediated βarr recruitment does not require the receptor’s C-terminus. This work reveals a new mode of GPCR activity regulation.

### Structure determination of atazanavir-mediated megacomplex

As described in the companion paper, we used a NanoBiT-based assay to measure the co-localization of miniG-protein and βarr, and the results suggested the existence of an atazanavir-mediated GPCR-Gs-βarr megacomplex. However, capturing the structure of such a megacomplex is technically challenging due to its intrinsic flexibility and transient nature, as well as the solubility limit of atazanavir. Furthermore, the expression level of GPR119 is low, which further hinders the structural studies. Luckily, atazanavir functions as a pan-PAM for a group of GPCRs, including the β_2_AR, which is more readily expressed than GPR119. To maximize the likelihood of capturing the atazanavir stabilized megacomplex, we introduced three mutations to the β_2_AR (K^6.35^R, G^6.38^S and I^6.39^V) to mimic the atazanavir binding pocket in GPR119. The resulted mutant receptor, β_2_AR-3mut, exhibits 2-fold higher of the maximum effect (E_max_) for atazanavir compared to β_2_AR_WT. The β_2_AR-3mut was then chosen for further investigation of the structure of megacomplex **(Fig.1a).**

Despite various stabilization strategies were tested, including NanoBiT-tethering strategy^20^ and receptor-miniGs fusion strategy^21^, the GPCR-G protein-βarr megacomplex remained unstable, and the density of βarr1 was not distinguishable in 2D averages. To further improve the stability of βarr1, a receptor-βarr1 fusion strategy was later tested by fusing the constitutively active human βarr-1(1-393) to the C terminus of receptor, a similar method has been used to obtain several high-resolution structures of the rhodopsin-βarr complex^6^ and the 5-HT_2B_-βarr complex^22^. Additionally, scFv30 was directly fused to the C-terminus of βarr to improve the stability of βarr1^22^ (**Fig. 1a**). The purified receptor-βarr1-scFv30 fusion protein was then mixed with heterotrimeric Gs protein to form the megacomplex, which was further purified by size exclusion chromatography (SEC) (Extended Data Fig. 1, see methods for detail). An ultra-potent β_2_AR agonist, BI-167,107, was used as the orthosteric ligand to stabilize the active conformation of the receptor^5,23^. Nb35, a nanobody frequently used to stabilize GPCR-Gs protein complexes^5^, was excluded from the cryo-EM sample preparation with the concern that it might compete with βarr1.

**Fig. 1.**
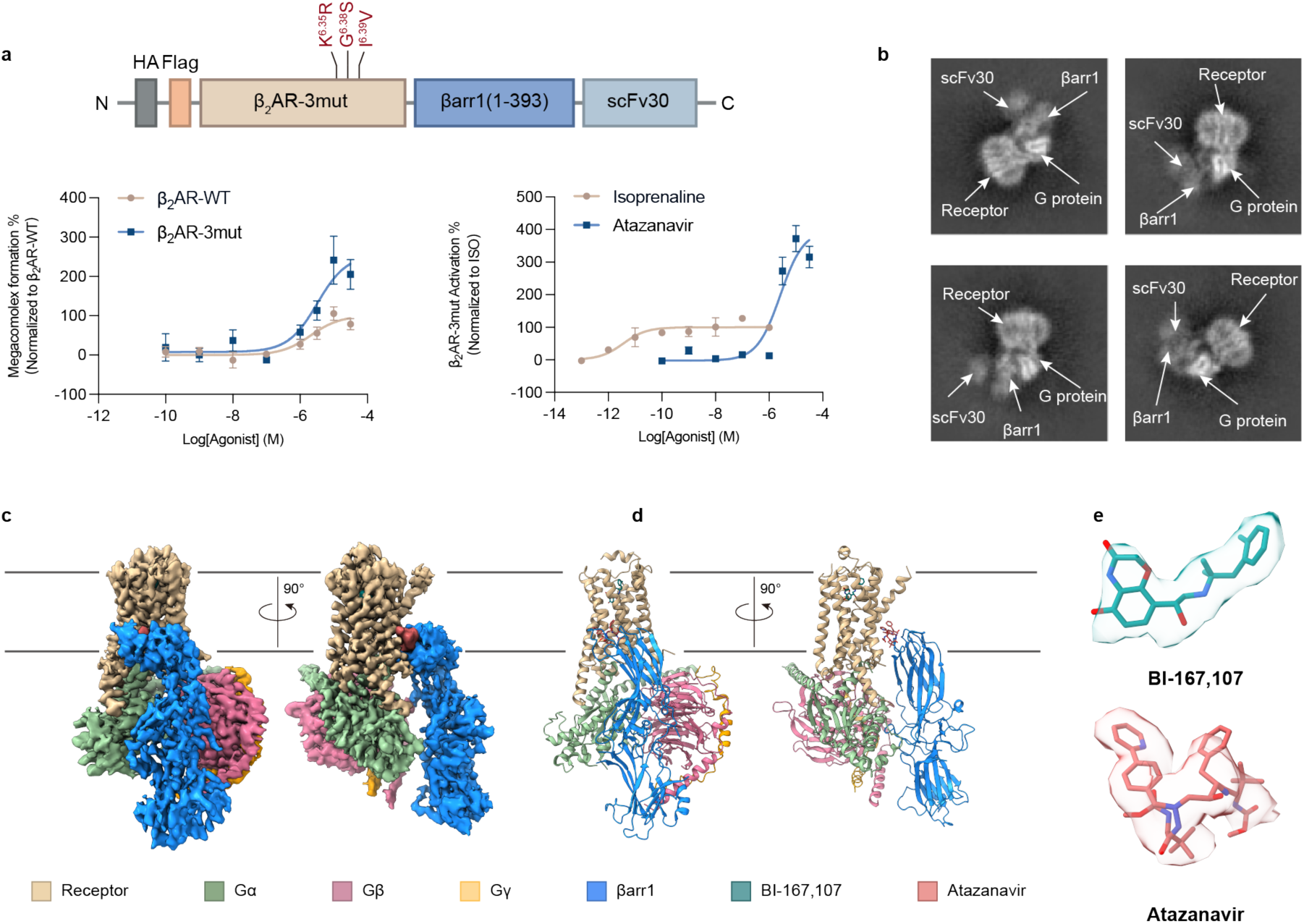
Structure determination of atazanavir-stabilized megacomplex. **a,** Construct used for the determination of megacomplex. The truncated human βarr1 (1-393) was directed fused to the C terminus of receptor, while a scFv30 was fused to the C terminus of βarr1 to facilitate stability. Three mutations (K^6.35^R, G^6.38^S and I^6.39^V) were introduced on the β_2_AR to mimic the binding pocket of atazanavir in GPR119, resulting in a chimeric receptor named as β_2_AR-3mut. The β_2_AR-3mut behaves better on both atazanavir-induced megacomplex formation and Gs protein activation. Data are shown as Mean ± S.E.M. of 3 independent replicates. **b,** The 2D averages of the Cryo-EM sample depict each component of the megacomplex. **c-d,** Cryo-EM map (c) and model (d) of atazanavir-stabilized GPCR-G protein-βarr1 megacomplex. The position of membrane bilayer is predicted using OPM server (https://opm.phar.umich.edu/ppm_server) and indicated as gray lines. β_2_AR-3mut, tan; Gα_s_, dark green; Gβ, violet red; Gγ, orange; βarr1, blue. **e,** Cryo-EM densities and models of BI-167,107 (dark cyan) and atazanavir (dark red).

Initial cryo-EM image processing revealed a subset of particles representing the megacomplex, with the features of βarr1 visible on 2D averages. To determine the structure of megacomplex, a total of 28,545 cryo-EM images were collected and processed (Extended Data Fig.2 **and Extended Data Table.1**). After iterative classification, all components of the megacomplex became clearly visible on 2D averages **(Fig.1b).** However, due to the heterogeneity of the megacomplex, initial attempts to reconstruct the entire megacomplex only resulted in a low-resolution map, with the cryo-EM density of βarr1 portion appeared blurred compared to the receptor-G protein complex portion. Despite this, the low-resolution map indicated that βarr1 attached to the TM6/TM7 of the β_2_AR-3mut and resided in the groove between the G_s_⍺ and the Gβγ. Of note, our previously determined structures of GPR119/β_1_AR-atazanavir revealed that atazanavir binds to the pocket of receptor where βarr1 attached to, indicating the βarr might attach the receptor through atazanavir. Meanwhile, 3D variability analysis (3DVA)^24^ of the βarr1-scFv30 also reveals a range of conformations of βarr1, as it oscillated relative to the receptor-Gs complex (approximately 40° movement between Gα subunit and Gβγ subunit, **Extended Data** Fig. 3), which might have been the major issue in global reconstruction and refinement. To gain deeper insight into atazanavir’s role in the assembly of the megacomplex, 3D classifications and refinements focused on the receptor-atazanavir-βarr1 interface were iteratively performed to identify particles with a homogeneous conformation of βarr1 (Extended Data Fig.2, see methods for detail). Finally, a 3.2 Å resolution cryo-EM map was resolved. The electron densities are clear enough to support model building for most components of the megacomplex (**Fig. 1c**), except for the scFv30 portion. The high-quality density map also clearly defines the binding mode of BI-167,107 and atazanavir in the orthosteric pocket and the intracellular allosteric site, respectively (**Fig. 1d and Extended Data** Fig 4).

### Atazanavir-mediated assembly of megacomplex

The atazanavir induced GPCR-G protein-βarr1 megacomplex revealed a noncanonical ‘pendulum’ conformation of βarr1, which is structurally compatible with the heterotrimeric Gs protein (**Fig. 1c**). In this megacomplex structure, a part of C-lobe of βarr1 is embedded in a detergent micelle, possibly due to its direct attachment to atazanavir near the transmembrane region of receptor. Of note, the edge of βarr1 C-lobe was reported to attach to or penetrate into the lipid bilayer when βarr1 bound to GPCRs^25^. The finger loop of βarr1, which is generally inserted into the transmembrane core of receptors in other structures^6,13,16,17,22,26–29^, is instead exposed to the solvent and positioned near the Gα subunit (**Fig. 1c**). The detailed conformational state of βarr1 will be discussed later.

In the structure, atazanavir is sandwiched between the TM6/7 region of the receptor and the 197-loop (R188-L197) of βarr1 (**Fig.1c and Fig.2a-b**). The 2-phenylpyridine moiety of atazanavir is adjacent to the TM6 side of the receptor, staying in proximity to G^6.42^ and V^6.39^. Additionally, the carbamate(1) moiety can form polar interactions with several hydrophilic residues at the intracellular region of TM6, mainly including R^6.35^, S^6.38^ and T^6.43^(**Fig. 2a,c**). Besides, the benzene ring and connected carbamate(2) moiety engage with TM7 of receptor through F^7.48^, L^7.51^, I^7.52^ and R^7.55^. In agreement with the structure information, mutations of these residues impair the function of atazanavir-induced megacomplex formation at different levels (**Extended Data Fig.5a and c**). Notably, as the canonical orthosteric agonist, isoprenaline disfavors the formation of GPCR-G protein-βarr1 megacomplex and thus leads to the robust decrease of luminescence signal. (**Extended Data Fig.5b**).

**Fig. 2.**
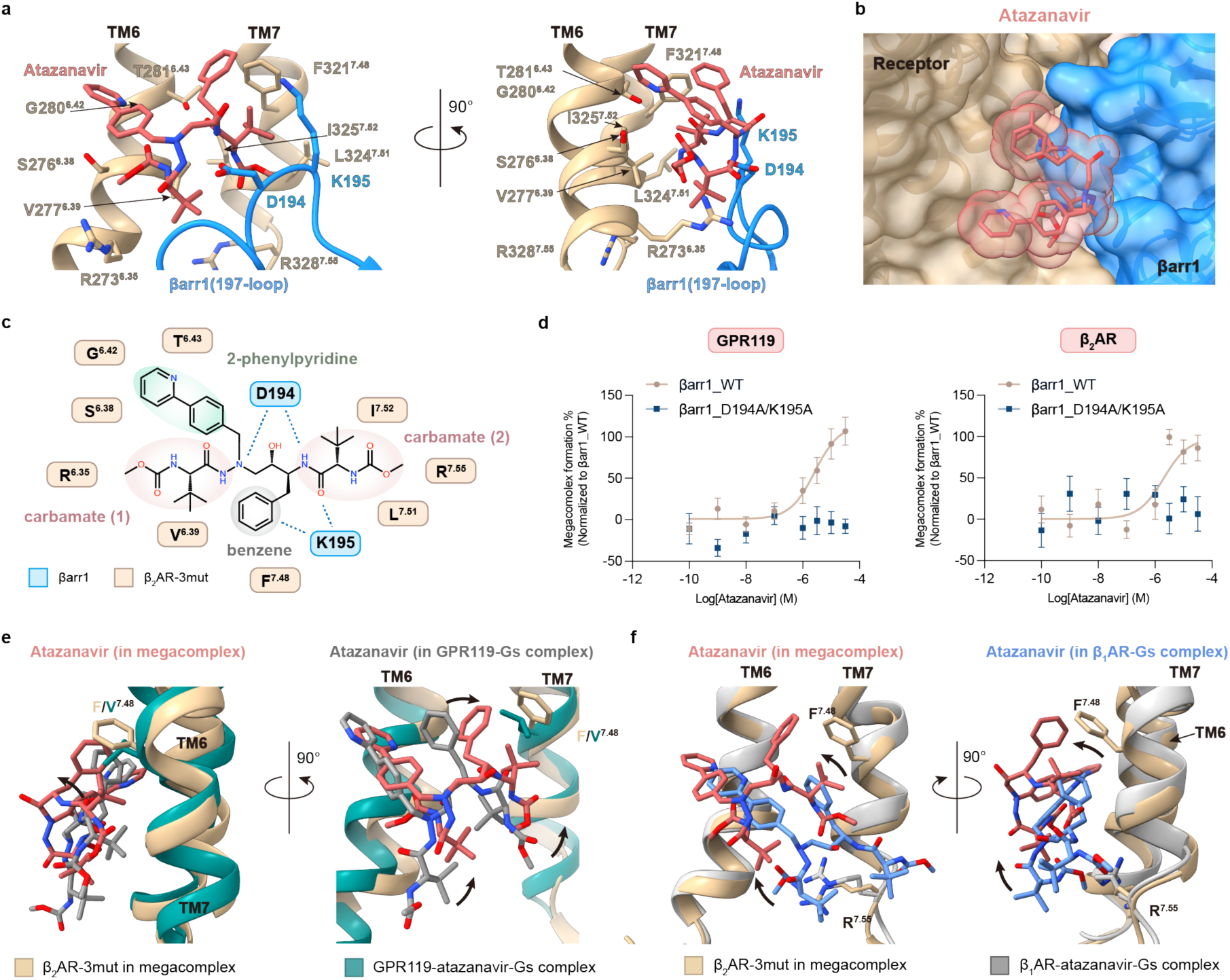
Atazanavir serves as a molecule glue to facilitate megacomplex formation. **a,** Detailed interactions at the interface of receptor-atazanavir-βarr1. The interface mainly includes the intracellular region of TM6/TM7 and the 197-loop of βarr1. The receptor portion is colored in tan and βarr1 is colored in blue. **b,** Atazanavir interacts with both the receptor portion and the βarr1 portion, occupying the cleft between receptor (tan) and βarr1 (blue) and serving as a molecular glue. **c,** Schematic showing the interface of receptor-atazanavir-βarr1, with different moieties indicated with distinct colors. **d,** The substitution of D194A/K195A disrupts the formation of megacomplex for both GPR119 (left) and β_2_AR (right). Data are shown as Mean ± S.E.M. of 3 independent replicates. **e-f,** The binding poses of atazanavir are slightly different in the megacomplex compared to the structure of GPR119-Gs (**e**) and β_1_AR-Gs (**f**). Conformational changes of atazanavir are indicated by arrows.

The structure of megacomplex also elucidates how βarr1 engages in the pendulum conformation. On the βarr1 side, the 197-loop remains close to atazanavir and thus takes account of the attachment of βarr1 to atazanavir (**Fig. 2a, c**). Specifically, D194 and K195 engage directly with atazanavir. D194 orients toward the central core of atazanavir, forming polar interactions with the compound, while K195 is situated near the benzene ring and a carbonyl group of atazanavir, contributing to the atazanavir-βarr1 recognition (**Fig. 2a, c**). As expected, the D194A/K195A mutation efficiently disrupts the formation of megacomplex for both GPR119 and β_2_AR (**Fig.2d**), without impacting the canonical recruitment of βarr1 to the receptor (Extended Data Fig. 6a-b). Meanwhile, we also employed a bioluminescence resonance energy transfer (BRET)-based co-localization assay (Extended Data Fig. 6c, see methods for detail) to evaluate the influence of D194A/K195A mutation of megacomplex assembly for β_2_AR, which results are consistent (Extended Data Fig. 6d**, left**). These structural insights, combined with our mutagenesis studies, shed light on the molecular basis of megacomplex assembly, with atazanavir functioning as a molecular glue to mediate the megacomplex formation (**Fig. 2b**).

In the companion manuscript, we reported the structures of atazanavir bound to the GPR119-Gs complex and the β_1_AR-Gs complex, providing key insights into the structural basis for atazanavir’s activation of these two distinct GPCRs. Structural comparison reveals that atazanavir binds within the same pocket near the intracellular ends of TM6 and TM7 across all three structures, despite subtle conformational changes. Overall, the binding pose of atazanavir in the megacomplex is more similar to its position in GPR119 than in β_1_AR (**Fig. 2e-f**). Furthermore, the association of βarr1 induces a slight repositioning of atazanavir, shifting it marginally away from the receptor interface (**Fig. 2e-f**).

The receptor portion of the megacomplex adopts a conformation nearly identical to that observed in the canonical GPCR-G protein complex^5,30^ (RMSD of Cα=0.84 Å, **Extended Data Fig.7a**), with TM6 displaying an outward-displaced conformation (**Extended Data Fig.7a**). The orthosteric agonist BI-167,107 also maintains a similar binding pose (**Extended Data Fig.7b**). Structure superimposition of the β_2_AR-Gs-Nb35-BI-167,107 and the megacomplex indicates that the Gα subunit retains a similar conformation, while the Gβγ subunit exhibits slight twists relative to its original position^5^ (**Extended Data Fig.7c**). Notably, we did not add Nb35 during the structural study of the megacomplex. The altered positioning of the Gβγ subunit may be attributed either to the presence of βarr1 or to the absence of Nb35.

### Active conformation of βarr1

The structure of atazanavir-mediated megacomplex highlights the active conformation of βarr1. Compared to its inactive structure^31^ (PDB ID: 1G4M), the C lobe of βarr1 in the megacomplex structure exhibits an interdomain twist relative to the N lobe. This interdomain conformational change, observed in many GPCR-βarr1 complex^6,14–17,22,26–29^, is considered as a hallmark of βarr1 activation (**Fig. 3a**).

**Fig. 3.**
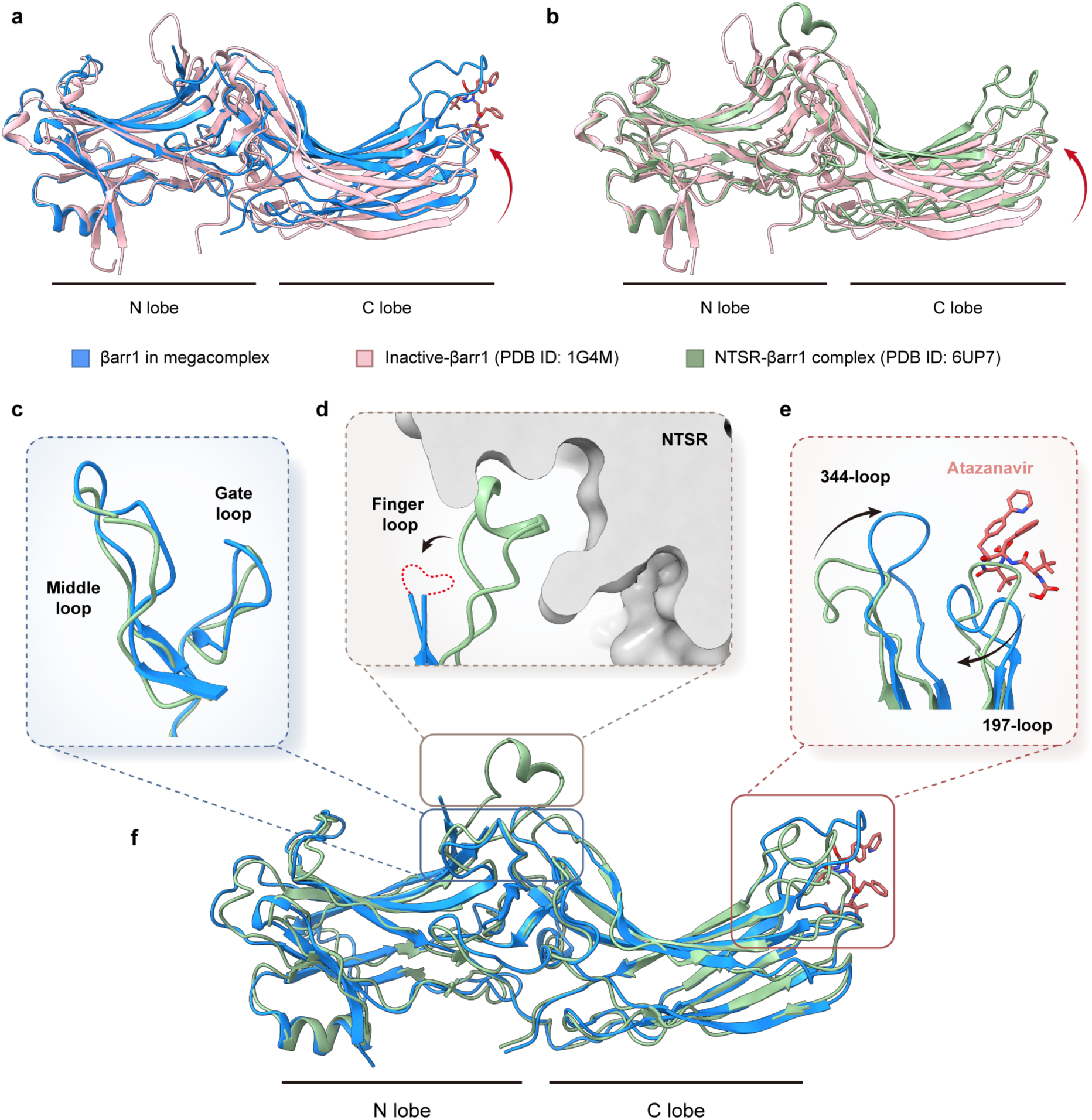
The active conformation of βarr1 in the megacomplex structure. **a-b,** Overlay of βarr1 in the megacomplex structure (blue, **a**) or NTSR1-bound βarr1 (PDB ID: 6UP7, dark green, **b**) with inactive βarr1 (PDB ID: 1G4M, pink). Compared to the inactive βarr1, the C lobe of βarr1 undergoes an interdomain twist in both structures, as indicated by the red arrow. **c-f,** Overlay of βarr1 in the megacomplex structure (blue) with NTSR1-bound βarr1 (PDB ID: 6UP7, dark green, **f**). The middle loop and gate loop adopt similar conformations (**c**). The finger loop of βarr1 is invisible in the megacomplex structure, probably due to the flexibility and lacking direct interaction with receptor. This loop usually turns into a helix in other core-engagement structures (**d**). Binding to atazanavir induces conformational changes of the 197-loop and the 344-loop of βarr1 in the megacomplex structure, as indicated by the arrows (**e**).

The βarr1 portion of megacomplex maintains an overall conformational state similar to that reported in the NTSR-βarr1^32^ structure (RMSD of Cα=1.19 Å), with notable differences primarily in the central crest and C-edge loops (**Fig. 3f**). Specifically, while the middle and gate loops within the central crest adopt conformations consistent with those in the NTSR-βarr1^32^ structure (**Fig. 3c**), the finger loop exhibit a distinct state. Typically, during core engagement, the finger loop inserts into the transmembrane bundle; however, in the megacomplex, the cryo-EM density of this loop is blurred and insufficient for model building, likely due to a lack of direct contacts with other components (**Fig. 3d**). This flexibility is further exacerbated by the dynamic nature of the entire βarr1 molecule (Extended Data Fig.3). Additionally, the attachment of atazanavir induces conformational changes in the 197-loop and 344-loop of βarr1. Compared to the NSTR-βarr1structure, the 197-loop shifts towards the 344-loop, forming a pocket that accommodates atazanavir binding. Concurrently, the 344-loop moves closer to the atazanavir binding site, potentially due to interactions between these elements (**Fig. 3e**).

### The non-canonical binding mode of βarr1 in megacomplex

Most reported GPCR-βarr complex structures capture the core-engagement conformation of the βarr. In these core engagement conformations, βarr adopts a range of orientations and tilted conformations. Structure alignment of the megacomplex with previously reported βarr1 complexes bound to NTSR1^26,28^, M_2_R^17^, β_1_AR^16^, V_2_R^15^, 5-HT_2B_R^22^, CB_1_R^27^ and mGlu3R^29^ reveals that the N lobe of βarr maintains a similar position relative to the receptors (**Fig.4a-b**). This consistency is due to that the majority of the interactions between the receptor and βarr are mediated through the N lobe, as the phosphorylated C-terminus of GPCRs typically inserts into the groove at the N lobe of βarr1. Conversely, the C lobe of βarr exhibits greater variability in its positioning and lack of direct interaction when bound to these different receptors (**Fig.4a-b**).

The binding mode of βarr1 in the megacomplex differs significantly from that observed in the core engagement structures. In the megacomplex, the predominant interactions originate from the C lobe of βarr1, with residues D194 and K195 (**Fig.2a, c**) directly interacting with atazanavir. As a result, the C lobe of βarr1 adopts a relatively stable conformation near the TM6/7 of receptor. Conversely, the N lobe of βarr1, lacking direct interactions with the receptor’s C-terminus and transmembrane core, exhibits greater flexibility (Extended Data Fig.3).

As mentioned earlier, two structures have captured the tail engagement conformation of βarr1. One of these structures, which represents the first megaplex structure, was determined using a β_2_V_2_R chimeric receptor (**Fig.4 c-d**). This chimeric receptor was constructed by replacing the C-terminus of the β_2_AR (a Class A GPCR) with that of the V_2_R (a Class B GPCR)^13^. In this structure, βarr1 interacts with the receptor exclusively through the phosphorylated C-terminus, leaving the intracellular pocket of the receptor accessible for G protein coupling and activation (**Fig.4d**). In contrast, the atazanavir-stabilized megacomplex exemplifies an allosteric agonist-mediated assembly of GPCR-G protein-βarr megacomplex, wherein βarr1 attaches to the TM6/7 region of the receptor through atazanavir-mediated interactions (**Fig.4d**). Despite the different mechanisms underlying the assembly of the GPCR-G protein-βarr1 megacomplex, both configurations ultimately result in sustained G protein signaling.

**Fig. 4.**
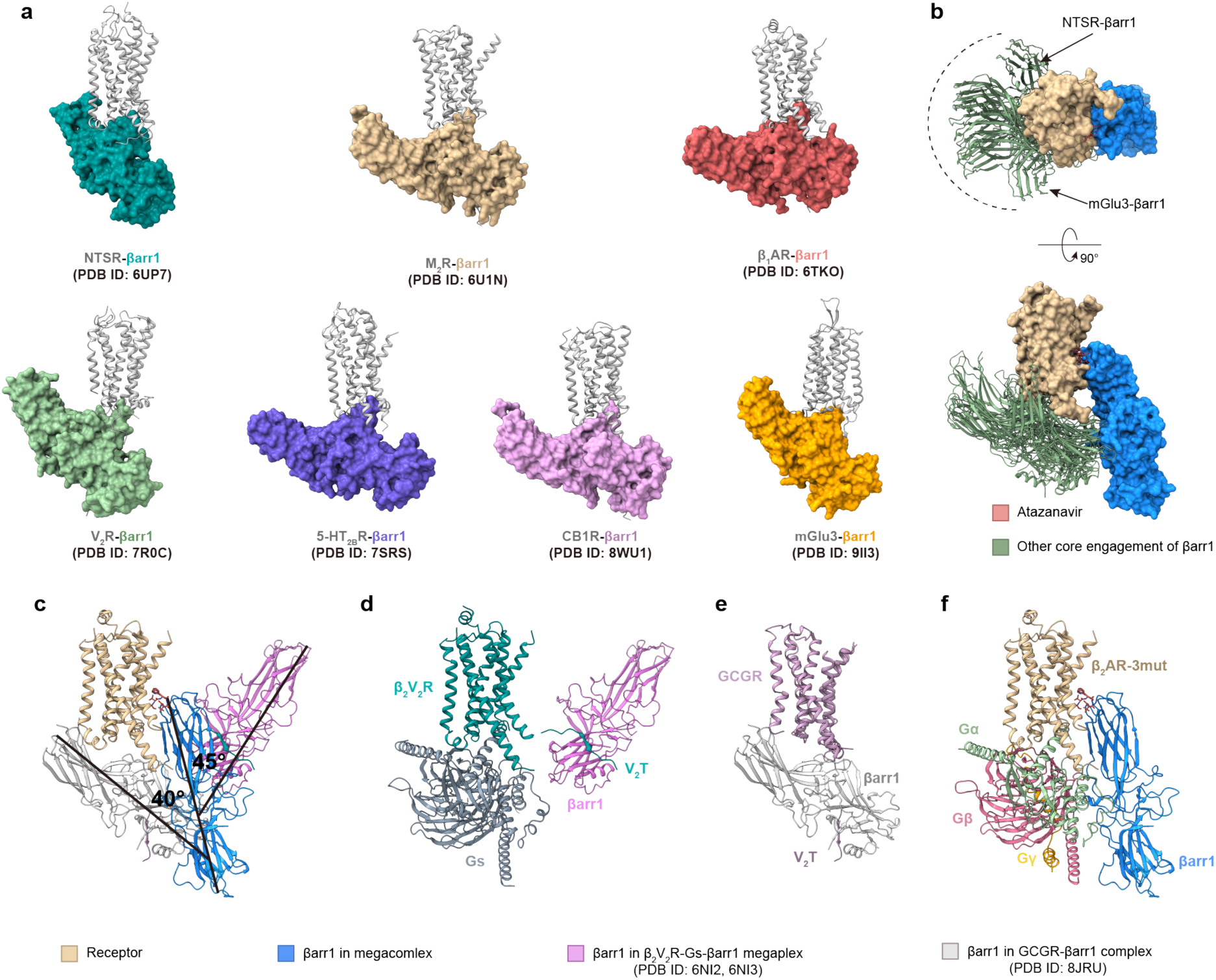
Non-canonical binding mode of βarr1 in megacomplex. **a-b,** Comparison of previous core engagement states of βarr1 in different GPCR complexes and in the atazanavir-induced megacomplex. In traditional core engagement states, βarr1 mainly interacts with receptor through its finger loop region and adopts various tilted conformations(**a**). Overlaying these core-engagement structures (colored in dark green) to the receptor (colored in tan) portion of the atazanavir-induced megacomplex suggests the non-canonical binding mode of βarr1 in the megacomplex (colored in blue, **b**). **c-f,** Structural comparison of atazanavir-stabilized megacomplex with β_2_V_2_R-Gs-βarr1 megaplex and GCGR-βarr1 complex. **(c)** For the structure of β_2_V_2_R-Gs-βarr1 megaplex, β_2_V_2_R-G_s_ subcomplex (PDB ID: 6NI3) and the βarr1-V_2_T subcomplex (PDB ID: 6NI2) are docked into the cryo-EM map of β_2_V_2_R-G_s_-βarr1 megaplex (EMDB-9377). The βarr1 in the β_2_V_2_R-Gs-βarr1 megaplex (plum) exhibits a 45° difference with the βarr1 in the structure of atazanavir-stabilized megacomplex (blue). The structure of GCGR-βarr1 (PDB ID: 8JRU) is aligned to the receptor (tan) portion of megacomplex, revealing a 40° difference between βarr1 in the megacomplex structure (blue) and βarr1 in the GCCR-βarr1 structure (silver). In the structure of β_2_V_2_R-Gs-βarr1 megaplex, the βarr1 (plum) mainly interacts with the receptor (dark green) through the V_2_T tail (dark green) and stays nearby the transmembrane region of receptor (**d)**. In the structure of GCGR-βarr1 complex, βarr1 stays in a tail-engagement mode near Helix 8 of GCGR (mauve). The C terminus of V_2_R (V_2_T) is also inserted into the groove of βarr1 (**e**). While for the atazanavir-induced megacomplex, the βarr1 engages with receptor in a different manner (**f**).

The other structure is exemplified by the GCGR-βarr1 complex^14^. In this structure, βarr1 primarily interacts with the Helix 8 of GCGR via its N lobe, while the C lobe remains in proximity to the ICL1 of GCGR (**Fig.4c**). Comparing the megacomplex in this study and the GCGR-βarr1 structures reveals that βarr1 adopts a vertical orientation in both, with the C lobe near the receptor, but with a distinct tilt angle difference of approximately 40° (**Fig.4c-d**). Notably, the Helix8 binding mode of βarr in the GCGR structure is compatible with the Gα subunit but may lead to partial clashes with the Gβγ subunit. This suggests that when βarr1 binds to the receptor simultaneously with heterotrimeric G protein, it may adopt an alternative conformation^14^.

Taken together, βarr1 exhibits a distinct binding mode in our structure of atazanavir-mediated megacomplex compared to previously reported core engagement or tail engagement conformations. In this structure, βarr1 attaches to the receptor primarily through its C lobe and resides in the groove between Gα and Gβγ subunit, aligning compatibly with the heterotrimeric G protein. The α-helical domain (AHD) of the G protein undergoes large conformational changes when G protein bound to different nucleotides **(Extended Data Fig.8a)**. Structure alignments of the megacomplex with G proteins in various nucleotide-binding states – nucleotide-free^5^ (PDB ID: 3SN6, **Extended Data Fig.8b**), GDP-bound^33^ (PDB ID: 6EG8, **Extended Data Fig.8c**) and GTP-bound^34^ (PDB ID: 8UO3, **Extended Data Fig.8d**) – suggest that the conformation of βarr1 in the megacomplex is compatible with each of these states. This compatibility indicates the potential of this megacomplex to support sustained G protein activation.

### The formation of megacomplex is independent of the C-terminus of GPCR

Previous researches indicate that βarr-GPCR interactions primarily rely on the phosphorylation of the GPCR C-terminus, which typically inserts into a groove on βarr^6,13–17,22,26–29^. However, after resolving the cryo-EM structure of the megacomplex, we observed no extra electron density for the receptor’s C-terminus in the groove of βarr1 (**Fig. 5a**, note that the auto-inhibitory C-terminus of βarr1 was truncated). The absence of this electron density in our structure suggests that the formation of the megacomplex does not require a stable interaction between the βarr1 and the C-terminus of GPCR. To investigate the influence of the GPCR C-terminus on megacomplex formation, we engineered C-terminus truncated versions for both GPR119 and β_2_AR (termed GPR119_ Δ301 and β_2_AR_ Δ342, **Fig. 5b-c**). A NanoBiT-based colocalization assay, as well as a BRET-based colocalization assay, revealed that both mutants are still capable of forming the megacomplex under the stimulation of atazanavir (**Fig. 5d and e, left, Extended Data** Fig. 6d**, right**), confirming that the atazanavir-induced megacomplex assembly is independent of the GPCR C terminus. Moreover, even in the absence of C-terminus, βarr1 recruitment to the receptor is still achievable under stimulation of atazanavir (**Fig. 5d and e, middle**), while for the orthosteric agonists of these two receptors, AR-231453 and isoprenaline, βarr1recruitment is abolished (**Fig. 5d and e, right**).

**Fig. 5.**
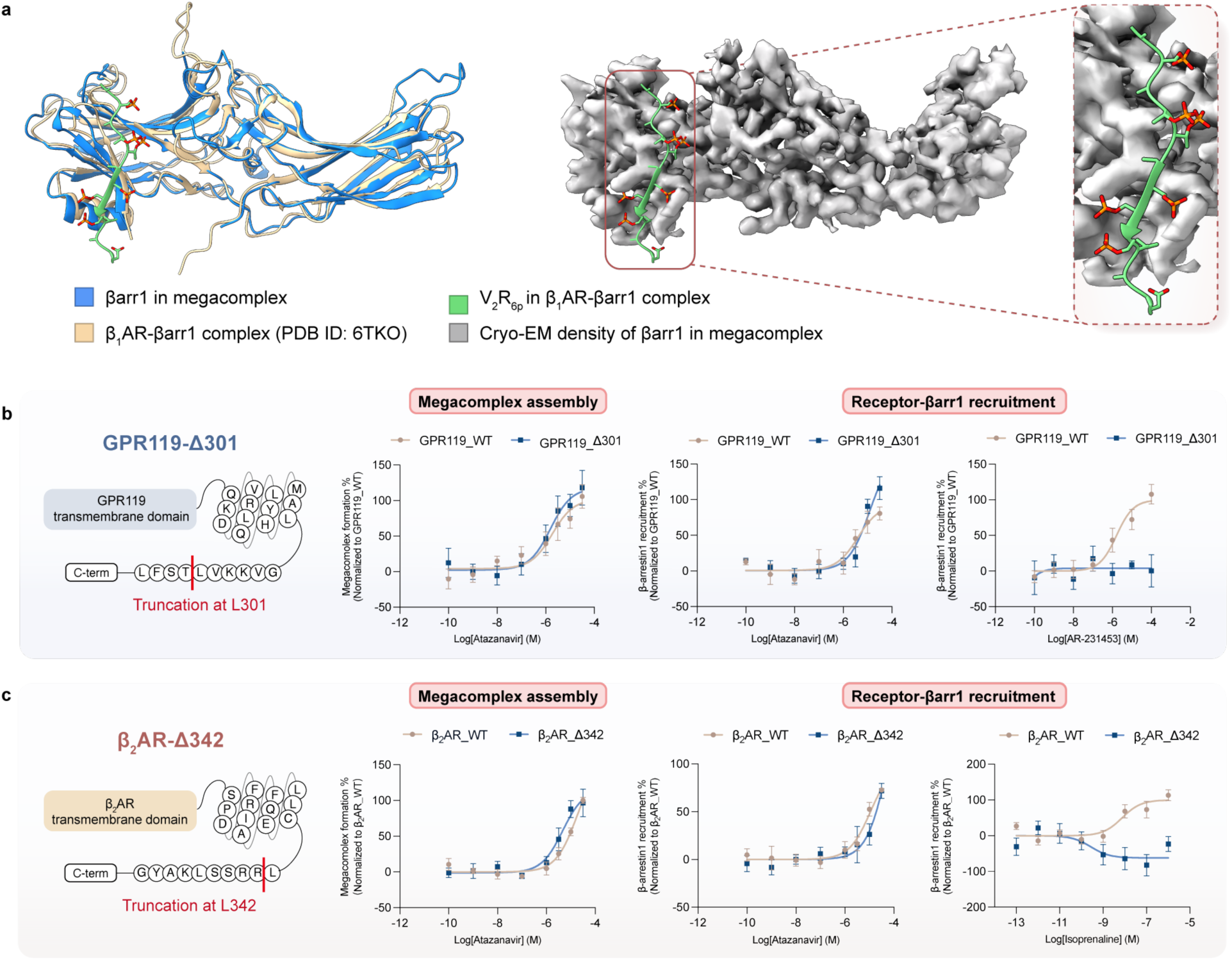
C-terminus-independent assembly of megacomplex **a,** Absence of GPCR C-terminus in the groove of βarr1 N lobe in the structure of megacomplex. Left, structure superimposition of βarr1 in the megacomplex (blue) with the counterpart in the β_1_AR-βarr1 complex (tan, PDB ID: 6TKO), the V_2_R_6P_ fused to β_1_AR (green) binds to the groove of βarr1 N lobe. Middle, the cryo-EM density (light gray) of βarr1 in megacomplex suggests the absence of GPCR C-terminus. Right, the zoom-in view of the GPCR C-terminus binding groove, no cryo-EM density (light gray) for the C-terminus of receptor is observed (green). **b-c,** Influence of C-terminus on megacomplex assembly and canonical βarr1 recruitment assays. The C terminus of GPR119 and β_2_AR are truncated at residue 301 (**c**, GPR119_Δ301) and residue 342 (**d**, β_2_AR_Δ342) respectively. Atazanavir induces the assembly of megacomplex as well as canonical βarr1 recruitment for GPR119_Δ301 (**c**) and β_2_AR_Δ342 (**d**), while the orthosteric agonist AR-231453 (**c**) and isoprenaline (**d**) cannot induce canonical βarr1 recruitment in the absence of GPCR C terminus. Data are shown as Mean ± S.E.M. of 3 independent replicates.

The association of βarr to the membrane is known as a key factor for GPCR-βarr1 interaction^25^. According to the megacomplex structure, the C-edge loop of βarr1 can penetrate into the membrane (Extended Data Fig. 9a). To investigate the role of membrane anchoring in megacomplex assembly, we introduced mutations into the C-edge loop of βarr1, impairing the membrane anchoring capability. Our results demonstrate the R331D/G332D/G333D mutation (βarr1_3D)^25^ abolishes the formation of megacomplex for both GPR119 and β_2_AR (Extended Data Fig. 9b), highlights that megacomplex formation also requires membrane anchoring of the C-lobe. Thus, membrane anchoring, together with the binding of 197-loop to atazanavir, drives the attachment of βarr1 to the receptor.

Membrane phosphoinositides (PIPs) binding to the C-lobe of βarr1 is crucial for the recognition between certain Class A GPCRs and βarr1. Disruption of the PIP_2_ binding site (K232/R236/K250) reduces βarr1 recruitment to the receptor^19^. Structure alignment revealed that the PIP_2_ binding site remains solvent-exposed in the megacomplex structure (**Extended Data Fig.9c**), indicating that megacomplex formation does not depend on PIP_2_ binding. To test this hypothesis, we constructed a PIP-binding-deficient mutant of βarr1 (K232Q/R236Q/K250Q, βarr1_3Q)^19^. The 3Q mutation did not affect megacomplex formation (**Extended Data Fig.9d**), confirming that the assembly of the megacomplex is independent of membrane phosphoinositide binding. Taken together, the assembly of atazanavir-induced megacomplex represents a non-canonical interaction mode, which is independent of both the GPCR C terminus and membrane phosphoinositides.

## Discussion

Given the critical role of GPCRs in mediating physiological and pathological responses, fine-tuning their function has garnered significant interest from both academic researchers and the pharmaceutical industry^35^. A variety of orthosteric ligands and allosteric modulators have been developed to address medical needs, or to deepen our understanding of GPCR modulation. The physiological functions of GPCRs rely on their interactions with downstream G proteins, as well as βarrs, which not only mediate the termination of G protein signaling, but also facilitate receptor endocytosis and other βarr-associated signaling pathways^1^. Advancing research has revised the traditional view that G proteins and βarrs are structurally exclusive. Studies have shown that some Class B GPCRs can form a stable tail engagement with βarr, while simultaneously activating G protein signaling, a phenomenon termed as the ‘megacomplex’^8,13^. However, this capability is facilitated by the inherently high affinity of Class B GPCRs for βarr. In contrast, assembly of such a megacomplex for Class A GPCRs is more challenging due to their low affinity to βarr^19^ (**Fig.6**, left).

**Fig. 6.**
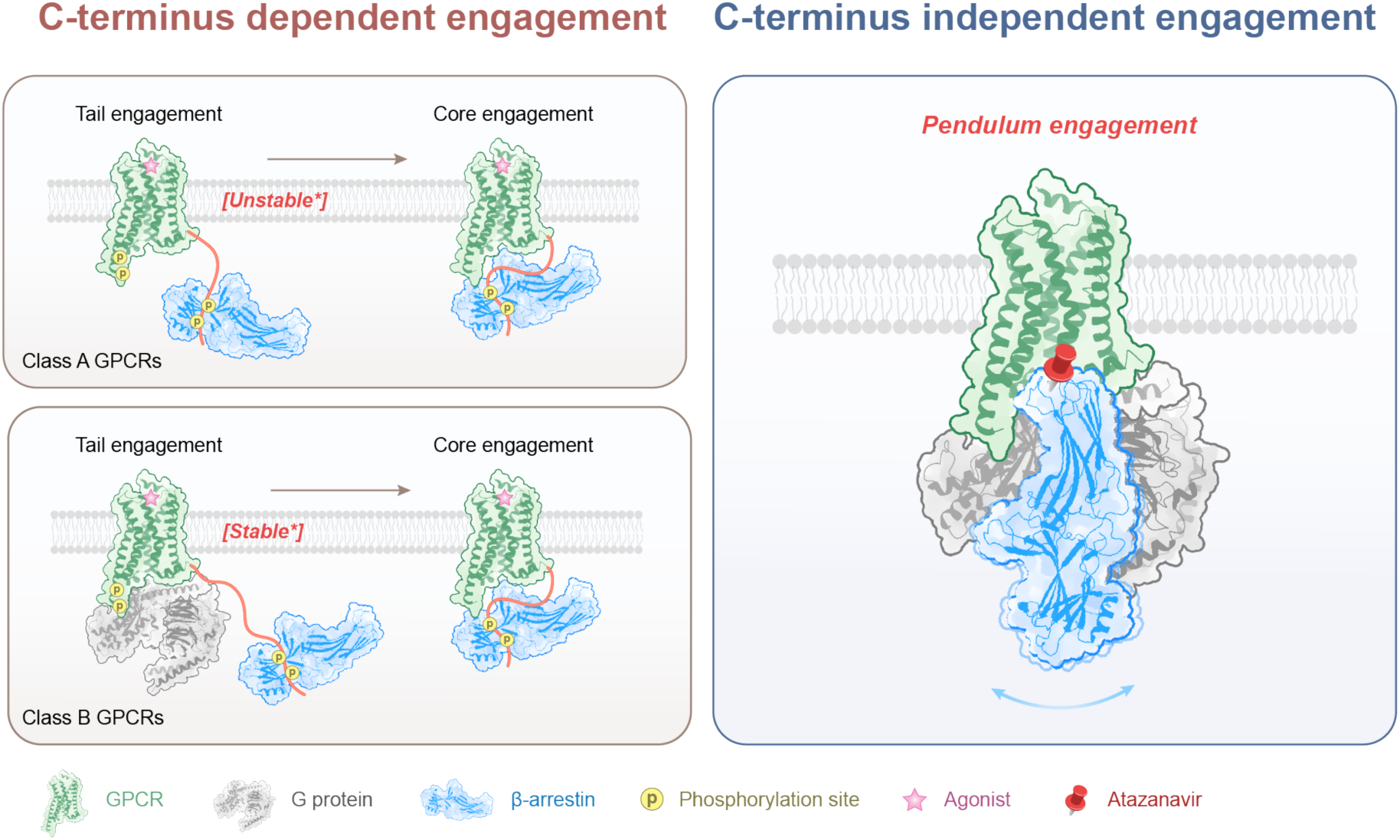
Atazanavir defines a novel mode of GPCR signaling modulation. This diagram elucidates different mode of GPCR-βarr interaction. **Left**, the canonical interaction between agonist-activated GPCRs and βarrs depends on the phosphorylated C-terminus of receptor, and βarrs engage with the receptor through both tail engagement and core engagement. ‘Class B’ GPCRs can form stable tail engagement with βarrs, leaving the intracellular pocket of receptor available for G protein coupling to form a megacomplex. **Right,** atazanavir-induced assembly of megacomplex, which is GPCR C-terminus independent. βarr1 adopts a pendulum engagement manner and resides at the groove of G protein.

In this work, we present the structure of atazanavir-induced GPCR-G protein-βarr1 megacomplex (**Fig.1c-d**) and elucidate the mechanism by which atazanavir acts as a molecular glue to facilitate the assembly of megacomplex (**Fig.2a-b**). Traditionally, the interaction between the phosphorylated C-terminus of GPCR and the βarr NTD has been considered crucial for both tail engagement and core engagement of βarr (**Fig.6, left** and Extended Data Fig.7). In our megacomplex structure, we demonstrate that in the presence of atazanavir, βarr1 binds to receptor in a noncanonical pendulum engagement mode specifically stabilized by atazanavir, which is independent of the GPCR’s C-terminus (**Fig.6, right**). This mechanism provides a new manner for Class A GPCRs to induce sustained G protein signaling, representing a new approach to GPCR function modulation.

The canonical paradigm of the competitive relationship between G protein and βarr has driven the development of biased agonists^36^. Unlike balanced agonists, which activate both pathways equally, biased agonists exhibit a preference for either G protein or βarr signaling, potentially reducing on-target side effects. Consequently, biased agonists are considered to have enhanced therapeutic potential in many contexts. For example, the therapeutic effects of many GPCR drugs are mediated through the G protein signaling pathways, such as the anti-asthma effects of β_2_AR agonists^37,38^ and the analgesic effects of μOR agonists^39,40^. In these scenarios, β-arrestin-mediated internalization and desensitization can diminish drug efficacy, leading to issues such as drug tolerance. To address these challenges, earlier studies attempted to develop biased agonists for β_2_AR or μOR that preferentially activate the G protein pathway over the β-arrestin pathway^36–39^. While this approach aims to reduce β-arrestin recruitment, it is challenging to completely eliminate it, and the bias is often relative rather than absolute. In this work, we propose that atazanavir represents a new strategy to modulate GPCR functions by making βarr structurally and functionally compatible with the G protein. This compatibility allows βarr to coexist with the G protein rather than competing with it. As a result, the atazanavir-mediated megacomplex does not induce desensitization at the molecular level, potentially offering therapeutic advantages in these applications. Moreover, previous research indicates that the formation of such a megacomplex can enhance the receptor’s G protein signaling^18^. The therapeutic potential of these megacomplexes warrants further exploration. Our megacomplex structure provides detailed insights into the interaction among the GPCR, atazanavir, and βarr. This information can guide the development of novel allosteric modulators with similar activities of atazanavir, potentially leading to new therapeutic strategies.

During the cryo-EM data processing, we observed a small fraction of particles that appear to represent the GPCR-G protein-βarr1 megacomplex state but lack clear density for atazanavir (**Extended Data Fig.2a**). These observations suggest the potential for GPCR-G protein-βarr1 megacomplex formation in the absence of atazanavir and raise the intriguing possibility that this pendulum engagement of βarr1 may represent an intrinsic binding mode between the receptor and βarr1, such as a pre-coupled state of βarr1 to the receptor. Atazanavir binding to the pocket may only strengthen this interaction. Notably, in the atazanavir-stabilized megacomplex structure, direct interactions between the receptor and βarr1 (defined by residues within 4 Å distance) could be observed (**Fig.2a**). However, we cannot rule out the possibility that the lack of atazanavir density in these particles is due to an average effect of multiple less-stable atazanavir binding poses. Further investigations are required to fully elucidate the mechanisms underlying this potential engagement mode.

## Methods

### Cloning and expression

The wild type human β_2_AR, containing an N-terminal HA signal sequence followed by a FLAG tag, was subcloned into pcDNA3.1 vector for expression in Expi293F cells. Based on the structures of GPR119-Gs-atazanavir and β_1_AR-Gs-atazanavir complexes described in the companion manuscript, three mutations (K^6.35^R, G^6.38^S and I^6.39^V) were introduced to the β_2_AR to mimic the atazanavir binding pocket in GPR119, and the engineered receptor was further named as β_2_AR-3mut. To improve the occupancy of βarr1, the truncated human β-arretin1(1-393) was fused to the C-terminus of β_2_AR-3mut. Additionally, an engineered scFv30 was directly fused to the C-terminus of βarr1 to improve its stability. The resulting fusion protein is denoted as β_2_AR-3mut-βarr1(1-393)-scFv30.

To express the fusion protein, the plasmid encoding β_2_AR-3mut-βarr1 (1-393)-scFv30 was transfected into Expi293F cells (ThermoFisher) through PEI (the plasmid to PEI ratio is 1:3) when the cell density reached approximately 2.5 million/ml. 60 hours after transfection, the cell pellet was collected and used for protein purification.

### β_2_AR-3mut-β-arretin1 (1-393)-scFv30 fusion protein purification

To purify the fusion protein, the cell pellet was firstly thawed in a lysis buffer containing 20 mM HEPES pH=7.5, 10 mM NaCl, 1 mM EDTA, 10 μM tris(2-carboxyethyl) phosphine (TCEP), 100 nM BI-167,107, 50 μM atazanavir, 20 µg/ml leupeptin and 160 µg/ml benzamidine. After stirring at room temperature for 30 minutes, the membrane was further solubilized using a buffer containing 20 mM HEPES pH=7.5, 150 mM NaCl, 2 mM CaCl_2_, 2 mM MgCl_2_, 1% lauryl maltose neopentyl glycol (LMNG, Antrace), 0.1% cholesteryl hemisuccinate (CHS), 10 μM TCEP, 100 nM BI-167,107, 50 μM atazanavir, 20 µg/ml leupeptin and 160 µg/ml benzamidine and benzonase for 2 hours at 4 ℃. The solubilized receptors were separated from the insoluble debris by centrifugation at 18, 000 rpm for 30 minutes and loaded onto an M1 Anti-FLAG column (Sigma-Aldrich). The unwanted contaminations were removed by washing with 30 column volumes (CVs) of wash buffer containing 20 mM HEPES pH=7.5, 150 mM NaCl, 2 mM CaCl_2_, 2 mM MgCl_2_, 0.01% LMNG, 0.001% CHS, 10 μM TCEP, 100 nM BI-167,107 and 50 μM atazanavir. Then the protein was eluted from the affinity column using an elution buffer containing 20 mM HEPES pH=7.5, 150 mM NaCl, 0.01% LMNG, 0.001% CHS, 10 μM TCEP, 100 nM BI-167,107, 50 μM atazanavir, 5 mM EDTA and 0.2 mg/ml FLAG peptide. After that, the fusion protein was concentrated and further purified by size-exclusion chromatography (SEC) on a Superose 6 Increase 10/300 GL (Cytiva) equilibrated in SEC buffer containing 20 mM HEPES pH=7.5, 150 mM NaCl, 2 mM CaCl_2_, 2 mM MgCl_2_, 0.003% LMNG, 0.0003% CHS, 10 μM TCEP, 100 nM BI-167,107 and 50 μM atazanavir. The peak fractions were collected and concentrated for further megacomplex preparation.

### Gβ_1_γ_2_ expression and purification

The Gβ_1_γ_2_ was expressed and purified with a similar method as previously described^21^. Briefly, the *Trichoplusia ni* (Hi5) insect cells (Expression Systems) were transduced with the baculovirus encoding Gβ_1_γ_2_ when cell density was around 3 million/ml. 48 hours after transduction, the cell pellet was collected and resuspended into a lysis buffer containing 20 mM Tris-HCl pH=7.5, 10 mM NaCl, 5 mM β-mercaptoethanol (β-ME), 20 µg/ml leupeptin and 160 µg/ml benzamidine. After stirring at room temperature for 30 minutes, the membrane was collected and solubilized in a solubilization buffer containing 20 mM HEPES pH=7.5, 100 mM NaCl, 1% sodium cholate, 0.05% dodecyl maltoside (DDM, Antrace), 0.005% CHS, 5 mM β-ME, 20 mM imidazole pH=8.0, 20 µg/ml leupeptin and 160 µg/ml benzamidine for 2 hours at 4 ℃. The solubilized protein was separated from the insoluble debris by centrifugation at 18, 000 rpm for 30 minutes and incubated with Ni-NTA resin (Cytiva) for 2 hours at 4 ℃. After incubation, the resin was washed with 10 CVs solubilization buffer. After that, the detergent was slowly exchanged to 0.1% LMNG and 0.01% CHS. Finally, the Gβ_1_γ_2_ was eluted from the resin with the elution buffer containing 20 mM HEPES pH 7.5, 100 mM NaCl, 0.01% LMNG, 0.01% CHS, 300 mM imidazole pH=8.0 and 100 μM TCEP. Eluted Gβ_1_γ_2_ was pooled and dialyzed overnight in a dialysis buffer containing 20 mM HEPES pH 7.5, 100 mM NaCl, 0.01% LMNG, 0.001% CHS, 100 μM TCEP and 10 mM imidazole. 3C protease was added to the protein to cleave the 6×His tag. The second day, reverse Ni-NTA affinity chromatography was performed to further increase the purity. Purified Gβ_1_γ_2_ was concentrated, flash frozen in liquid N_2_ and stored at −80 ℃ for further use.

### Gα_s_ expression and purification

The Gα_s_ was expressed and purified with a similar method as previously described^33^. The bovine Gα_s_ protein with a N-terminal 6×His tag followed by a 3C protease site was subcloned into pET28a vector for expression in BL21(*DE3*) cells (Vazyme). Cells were grown in TB medium to OD_600_ of 1.5, then the protein expression was induced by addition of 1 mM Isopropyl β-D-1-thiogalactopyranoside (IPTG). After incubation at 22 ℃ overnight, the pellet was collected and resuspended in a lysis buffer containing 50 mM HEPES pH=7.5, 100 mM NaCl, 1 mM MgCl_2_, 10 μM Guanosine diphosphate (GDP), 5 mM β-ME, 20 mM imidazole pH=8.0, 20 µg/ml leupeptin and 160 µg/ml benzamidine. The cells were disrupted by sonication using a 60 % power, 3 second on / 3 second off cycles, 5 minutes. The supernatant containing Gα protein was separated from the cell debris by centrifugation at 18,000 rpm for 30 minutes and incubated with Ni-NTA resin for 2 hours at 4 ℃. The resin was washed with 30 CVs lysis buffer to remove unwanted contaminations. Then the Gα_s_ was eluted from the resin with the lysis buffer containing 250 mM imidazole pH=8.0. Eluted Gα_s_ was pooled and dialyzed overnight in dialysis buffer containing 20 mM HEPES pH 7.5, 100 mM NaCl, 100 μM TCEP, 1 mM MgCl_2_, 20 μM GDP and 10 mM imidazole. 3C protease was added to the protein to cleave the 6×His tag. The second day, reverse Ni-NTA affinity chromatography was performed to remove contaminations. Finally, the Gα was concentrated and further purified by SEC on Superdex 75 increase 10/300 GL equilibrated in SEC buffer containing 20 mM HEPES pH 7.5, 100 mM NaCl, 100 μM TCEP, 1 mM MgCl2, 20 μM GDP. The peak fractions were collected, concentrated, flash frozen in liquid N_2_ and stored at −80 ℃ for further use.

### Formation of megacomplex

The purified β_2_AR-3mut-β-arretin1 (1-393)-scFv30 fusion protein was incubated with 1.2 molar excess Gα_s_ and Gβ_1_γ_2_ protein in an incubation buffer containing 20 mM HEPES pH=7.5, 150 mM NaCl, 4 mM CaCl_2_, 2 mM MgCl_2_, 0.003% MNG, 0.0003% CHS, 50 μM BI-167,107, 100 μM atazanavir and 20 μM C_8_-PIP_2_. After incubation at room temperature for 1 hour, Apyrase (New England Biology) was added to hydrolyze remaining GDP. After another 1 hour, the complex was further purified by SEC on a Superose 6 Increase 10/300 GL equilibrated in SEC buffer containing 20 mM HEPES pH=7.5, 150 mM NaCl, 2 mM CaCl_2_, 2 mM MgCl_2_, 0.002% LMNG, 0.0002% CHS, 10 μM TCEP, 100 nM BI-167,107 and 50 μM atazanavir. The peak fractions were collected and concentrated for cryo-EM sample preparation.

### Cryo-EM sample preparation and data collection

The protein sample of megacomplex was applied to glow-discharged grids (Quantifoil Au 1.2/1.3 300 mesh). The blotted grids (blot time 4.5 second, blot force 0, at 8℃, 100 % humidity) were rapidly frozen in liquid ethane by using Vitrobot (FEI Mark IV, Thermo Fisher Scientific). The data collection was conducted on a 300 kV Titan Krios. The movies were captured with a bin2 pixel size of 1.08 Å using EPU data acquisition software in super resolution mode, with a defocus range of −1.1 to −1.6 μm. The total dose is about 50 e-/Å^2^ for each stack. Movies were collected and subjected to beam-induced motion correcting by using MotionCor2^41^ for further data process.

### Model building and refinement

For model building, the coordinate files of β_2_AR-Gs (PDB ID: 3SN6, use the receptor and G protein portion) and NTSR-βarr1 (PDB ID: 6UP7, use the βarr1 portion) were docked into the cryo-EM map of megacomplex using UCSF ChimeraX^43^, followed by iterative manual adjustment in COOT. Models and restrain files of atazanavir and BI-167,107 were generated by eLBOW^44^ in PHENIX^45^. The model of megacomplex was then automatically refined in PHENIX real space refinement and manually adjusted in COOT^46^. Final validation statistics used for Extend Table 1 were generated by PHENIX comprehensive validation (cryo-EM). All structure figures were prepared using UCSF ChimeraX.

### cAMP GloSensor assay

CHO-K1 cells were cultured in DMEM/F12 medium (Gibco) containing 10% FBS, at 37 ℃, 5% CO_2_. The day before transfection, CHO cells were seeded into 6-well plates in DMEM/F12 medium containing 10% FBS at the density of 2 ×10^5^ cells per ml. After overnight growth, the plasmids including receptor and 22F-pGloSensor (Promega) were co-transfected into CHO-K1 cells to detect the receptor activation-induced cAMP accumulation. After 24 hours, cells were digested by 0.05% trypsin, resuspended with CO_2_-independent medium (Gibco) containing 10% FBS and 2% GloSensor cAMP Reagent (Promega), and plated into 96-well plates at a volume of 90 μl. Cells were incubated at 37 ℃ for 1 hour followed by incubated at room temperature for 1 hour. After incubation, the basal luminescence of each well was determined and varying concentrations of compound were added into each well to detect luminescence increase. Luminescence signal of each well was measured every 30 seconds and data was processed by GraphPad Prism 9.0 (GraphPad LLC, CA) and shown as Mean ± SEM.

### NanoBiT-based megacomplex co-localization assay

The day before transfection, CHO-K1 cells were seeded into 6-well plates in DMEM/F12 medium containing 10% FBS at the density of 2 ×10^5^ cells per ml. After overnight growth, the plasmids including receptor, SmBiT-miniGs and LgBiT-βarr1 (or mutants) were co-transfected into CHO-K1 cells to detect the formation of megacomplex. 24 hours after transfection, cells were digested with 0.25% trypsin, resuspended with HBSS buffer containing 20 mM HEPES pH=7.5 and 10 μM coelenterazine and seeded into 96-well plates at a volume of 90 μl. After 2 hours incubation at room temperature, the basal luminescence of each well was determined and varying concentrations of compound were added into each well to detect luminescence increase. Luminescence signal of each well was measured every 30 seconds and data was processed by GraphPad Prism 9.0 (GraphPad LLC, CA) and shown as Mean ± SEM.

### BRET-based megacomplex co-localization assay

The day before transfection, CHO-K1 cells were seeded into 6-well plates in DMEM/F12 medium containing 10% FBS at the density of 2 ×10^5^ cells per ml. After overnight growth, the plasmids including receptor, *Renilla* luciferase (*R*Luc8)-miniGs and Venus-βarr1 (or mutants) were co-transfected into CHO-K1 cells to detect the formation of megacomplex. 24 hours after transfection, cells were digested with 0.25% trypsin, resuspended with DMEM/F12 medium, seeded into 96-well plate and incubated overnight. The next day, cell medium was replaced with HBSS buffer containing 20 mM HEPES pH=7.5 and incubated 15 min at room temperature. Varying concentrations of compound and 10 μM coelenterazine were added into each well and plates were detected for both luminescence at 485 nm and fluorescent emission at 535 nm. The ratio of Venus/Rluc was calculated per well, processed by GraphPad Prism 9.0 (GraphPad LLC, CA) and shown as Mean ± SEM.

### Receptor-βarr1 recruitment assay

The day before transfection, CHO-K1 cells were seeded into 6-well plates in DMEM/F12 medium containing 10% FBS at the density of 2 ×10^5^ cells per ml. After overnight growth, the plasmids including receptor-LgBiT (or mutants) and SmBiT-βarr1 (or mutants) were co-transfected into CHO-K1 cells to detect the recruitment between receptor and βarr1. 24 hours after transfection, cells were digested with 0.25% trypsin, resuspended with HBSS buffer containing 20 mM HEPES pH=7.5 and 10 μM coelenterazine and plated into 96-well plates at a volume of 90 μl. After 2 hours incubation at room temperature, the basal luminescence of each well was determined and varying concentrations of compound were added into each well to detect luminescence increase. Luminescence signal of each well was measured every 30 seconds and data was processed by GraphPad Prism 9.0 (GraphPad LLC, CA) and shown as Mean ± SEM.

### Expression level of receptor mutants

CHO-K1 cells were seeded in 12-well dishes at a density of 2 × 10^5^ cells per ml medium supplemented with 10% FBS, penicillin and streptomycin 1 day before transfection. For each well, receptor with different mutation or pcDNA3.1 (vehicle) was transfected to determine the expression level. After 1 day, cells are treated with 0.25% Trypsin and resuspended with HBSS containing 20 mM HEPES pH=7.5 to a volume is 300 μl. 10 nM AlexaFlour-488 labelled M1-anti FLAG antibody are mixed with cells for 15 min in dark. After incubation, cells are washed with HBSS containing 20 mM HEPES pH=7.5 for twice. Fluorescence of each sample is determined by Flow Cytometer at 488 wavelengths. Data was processed in FlowJo (BD Biosciences).

## Data availability

The cryo-EM density map of β_2_AR-3mut-Gs-βarr1-atazanavir has been deposited into the Electron Microscopy Data Bank (EMDB) under accession code EMDB-62885, the model coordinate of β_2_AR-3mut-Gs-βarr1-atazanavir has been deposited into the Protein Data Bank (PDB) under accession code 9LBL.

## Acknowledgements

We thank Prof. Brian K. Kobilka for discussion and suggestions, and Prof. Asuka Inoue for providing the plasmids for the NanoBiT assay. We thank the Tsinghua University Branch of China National Center for Protein Sciences (Beijing) as well as Shuimu Biosciences for supports with cryo-EM data collection. This work was supported by the Tsinghua-Peking Center for Life Sciences (X.L.), Beijing Frontier Research Center for Biological Structure, Tsinghua University (G.H., X.X. and X.L.), by National Natural Science Foundation of China (Grant 32122041 to X.L.) and by Tsinghua University Initiative Scientific Research Program (X.L.).

## Author contributions

X.L. conceived and supervised the project. G.H., Q.S., and X. L. designed the experiments. G.H., Q.S. and X.X. screened different strategies to prepare the megacomplex. G.H. and Q.S. purified the protein sample of receptor-G protein-βarr1 megacomplex with the help from X.S.. G.H., Q.S. and S.Z. prepared the cryo-EM sample and collected the data. G.H. and X.L. processed the cryo-EM data with the help and suggestions from F.K., S.Z. and C.Y.. G.H. and X.L. built the protein model. G.H. and Q.S. analyzed the model and performed the cellular functional assay. K.Y. synthesized the BI-167,107 under the supervision of X.C.. G.H, Q.S and X.L. wrote the manuscript with the inputs from all the authors.

## Materials & Correspondence

Correspondence and material requests should be addressed to Xiangyu Liu.

---

**Extended Data Fig. 1.**
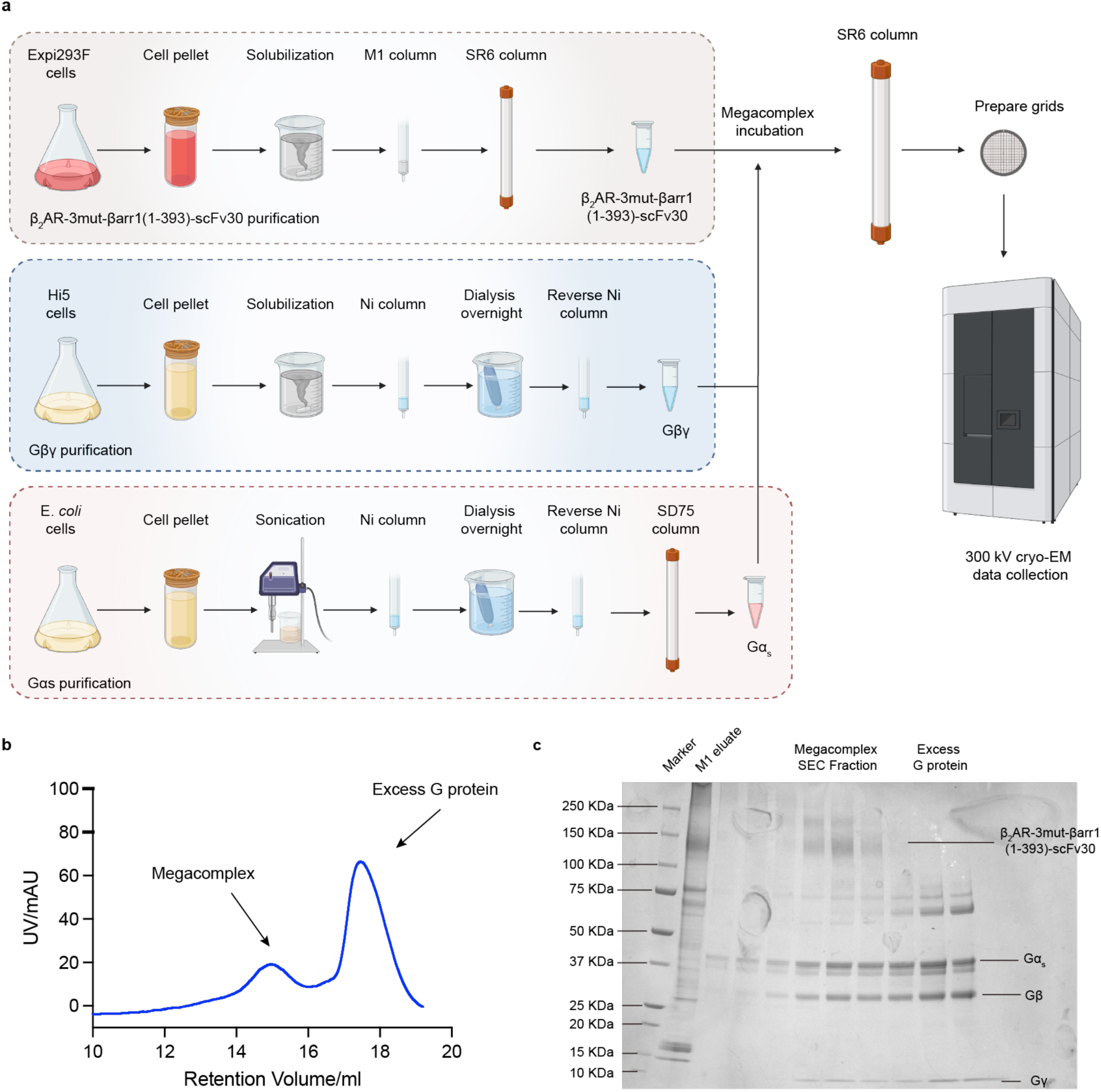
Atazanavir-stabilized megacomplex preparation. **a,** Flow diagram of protein expression, purification and megacomplex assembly. β_2_AR-3mut-βarr1(1-393)-scFv30, Gα_s_ and Gβ_1_γ_2_ were separately purified. Purified proteins were incubated together to form the megacomplex, excess G proteins were removed by size-exclusion chromatography (SEC). Peak fractions representing the megacomplex were collected for cryo-EM sample preparation and data collection. The figure was created with BioRender. **b,** The SEC profile of atazanavir-stabilized megacomplex and excess G proteins. **c,** SDS-PAGE gel of the protein samples from the final SEC.

**Extended Data Fig. 3.**
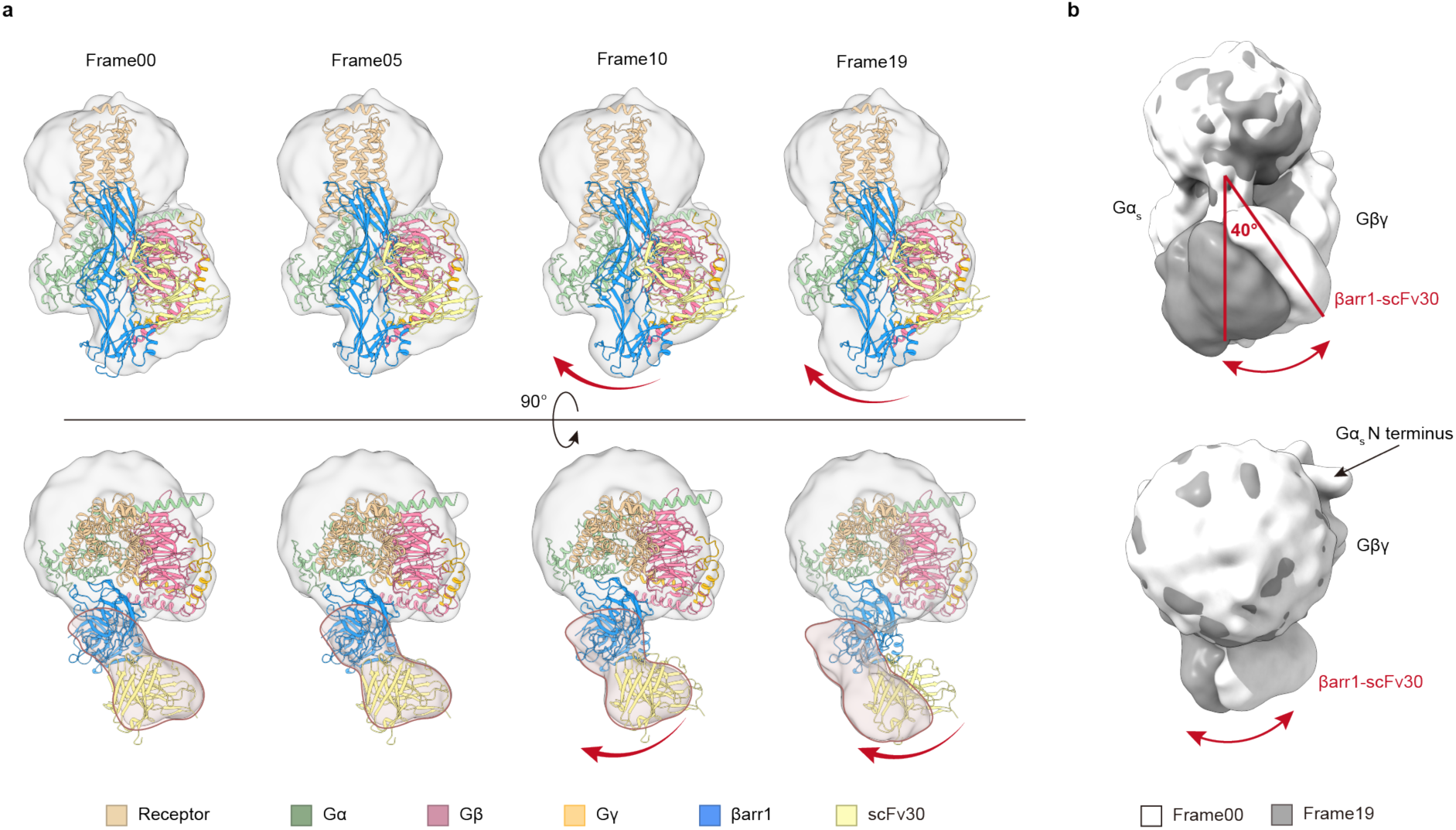
3D variability analysis (3DVA) on the heterogeneity of megacomplex a, Four different frames generated from the 3DVA suggest the movement of βarr1-scFv30 portion of the megacomplex relative to the receptor-Gs portion. The model of megacomplex is docked into the cryo-EM map of frame00. The movement is indicated by the red arrow, while red lines indicate the outline of βarr1-scFv30 cryo-EM map. **b,** βarr1-scFv30 portion oscillated approximately 40° movement between Gα subunit and Gβγ subunit, revealed by the cryo-EM map of frame00 (white) and frame19 (dark gray).

**Extended Data Fig. 4.**
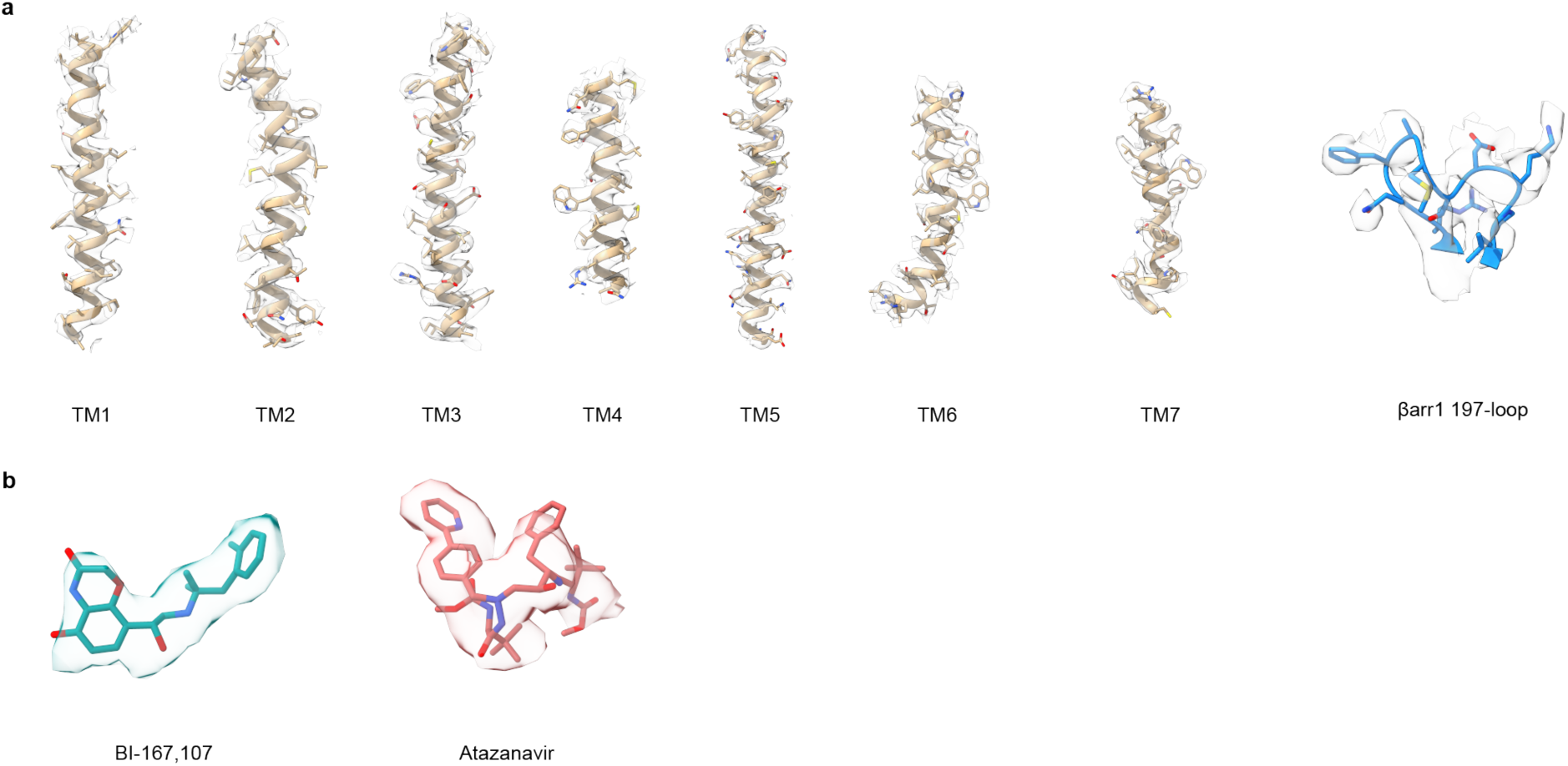
Cryo-EM densities of atazanavir-induced megacomplex **a,** Cryo-EM densities of transmembrane helices of β_2_AR-3mut and 197-loop of βarr1. **b,** Cryo-EM densities of BI-167,107 and atazanavir.

**Extended Data Fig. 5.**
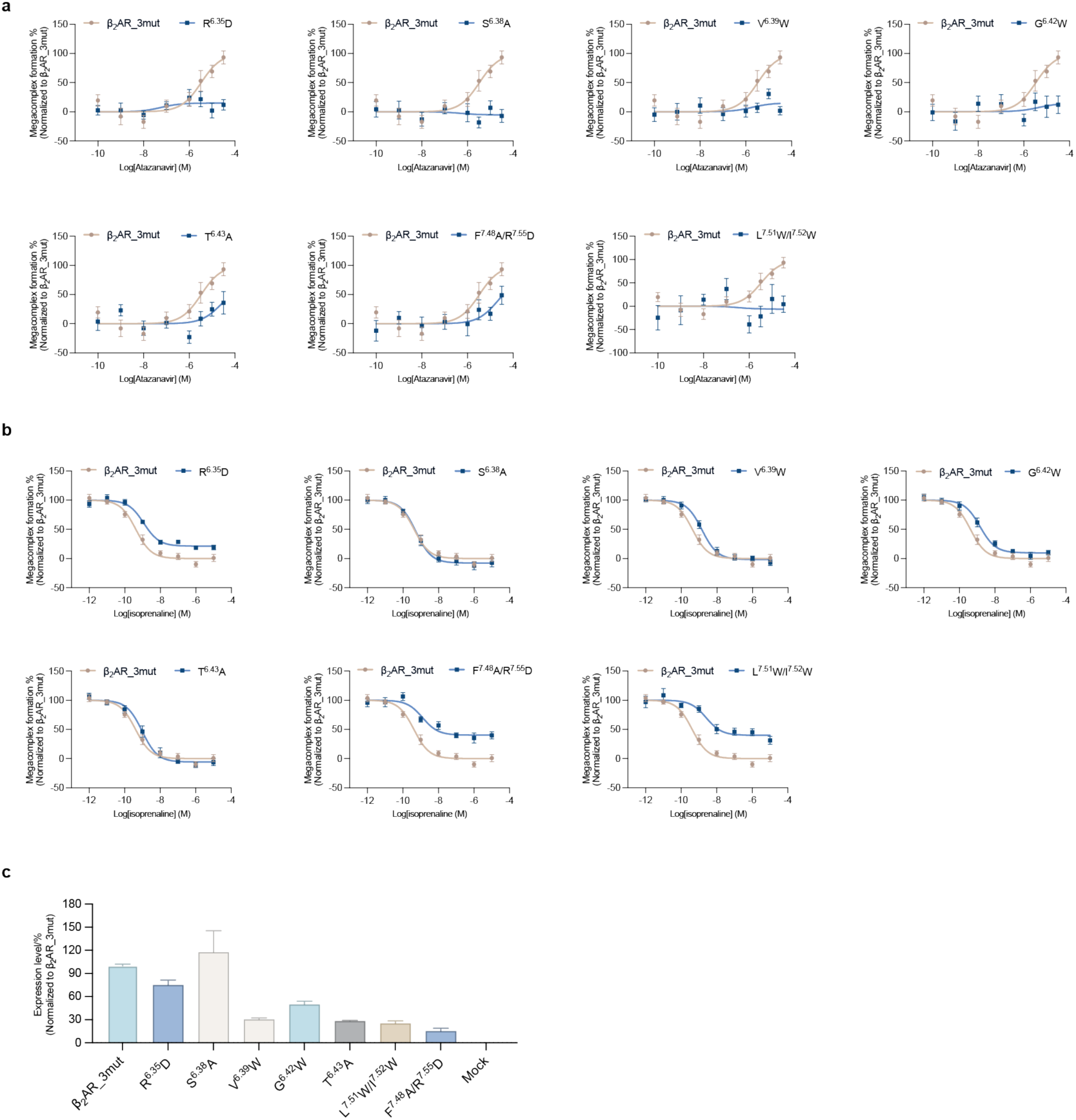
Mutations on β_2_AR-3mut influence megacomplex formation **a,** Mutations on the binding pocket of atazanavir influence the formation of megacomplex at different levels. **b,** Isoprenaline induce robust decrease of luminescence signal for all mutants. **c**, Relative expression level of each mutant and β_2_AR-3mut. Data are shown as Mean ± S.E.M. of 3 independent replicates.

**Extended Data Fig. 6.**
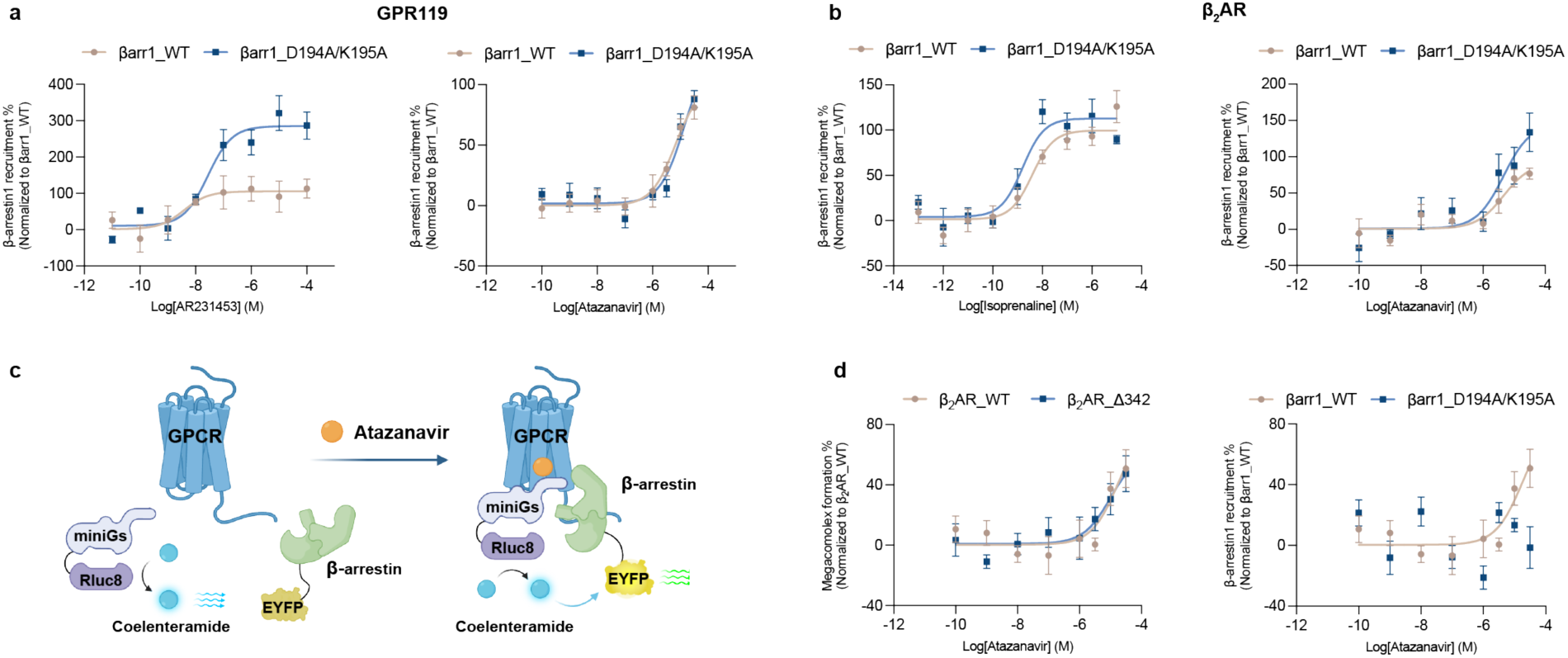
Functional assay on the βarr1 recruitment and megacomplex formation a-b, The βarr1_D194A/K195A can be recruited to both GPR119 (**a**) and β_2_AR (**b**) upon activation of orthosteric agonist (AR231453 for GPR119 and isoprenaline for β_2_AR) and atazanavir. **c,** Schematic of BRET-based miniGs and βarr1 colocalization assay. The Rluc is fused to the N terminus of miniGs and the Venus is fused to the N terminus of βarr1. Colocalization of miniGs and βarr1 leads to an increased BERT ratio. **d,** BRET-based colocalization assay also suggests the assembly of megacomplex is independent of receptor ‘s C terminus (**left**), and the D194/K195 is crucial for the assembly of megacomplex (**righ**t). Data are shown as Mean ± S.E.M. of 3 independent replicates.

**Extended Data Fig. 7.**
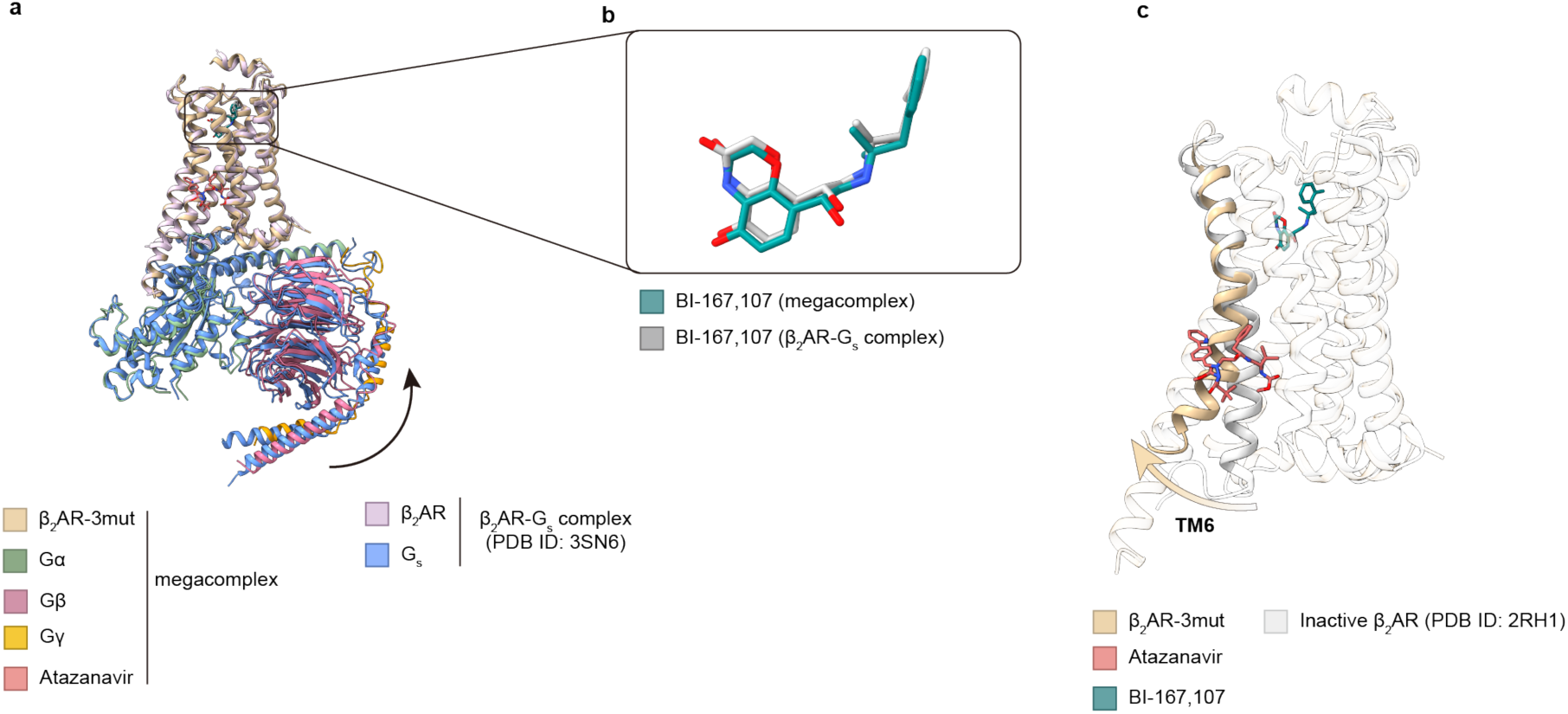
Structure alignment of the receptor-Gs portion of megacomplex with other β_2_AR structures. **a,** The β_2_AR-3mut (tan) adopts a similar conformation as it in the β_2_AR-Gs complex (PDB ID: 3SN6, mauve). The Gα_s_ (dark green) also stays in a similar conformation, while the Gβγ (violet red and orange) subunit twist slightly, indicated by the black arrow. The G_s_ portion of β_2_AR-Gs complex is colored in dodger blue. **b,** The orthosteric agonist BI-167,107 (dark cyan) adopts a similar pose as it in β_2_AR-Gs complex (PDB ID:3SN6, BI-167,107 is colored in silver) **c,** The receptor stays in the active state, with the TM6 adopts an outward movement compared to the inactive state of β_2_AR (PDB ID: 2RH1, colored in light gray).

**Extended Data Fig. 8.**
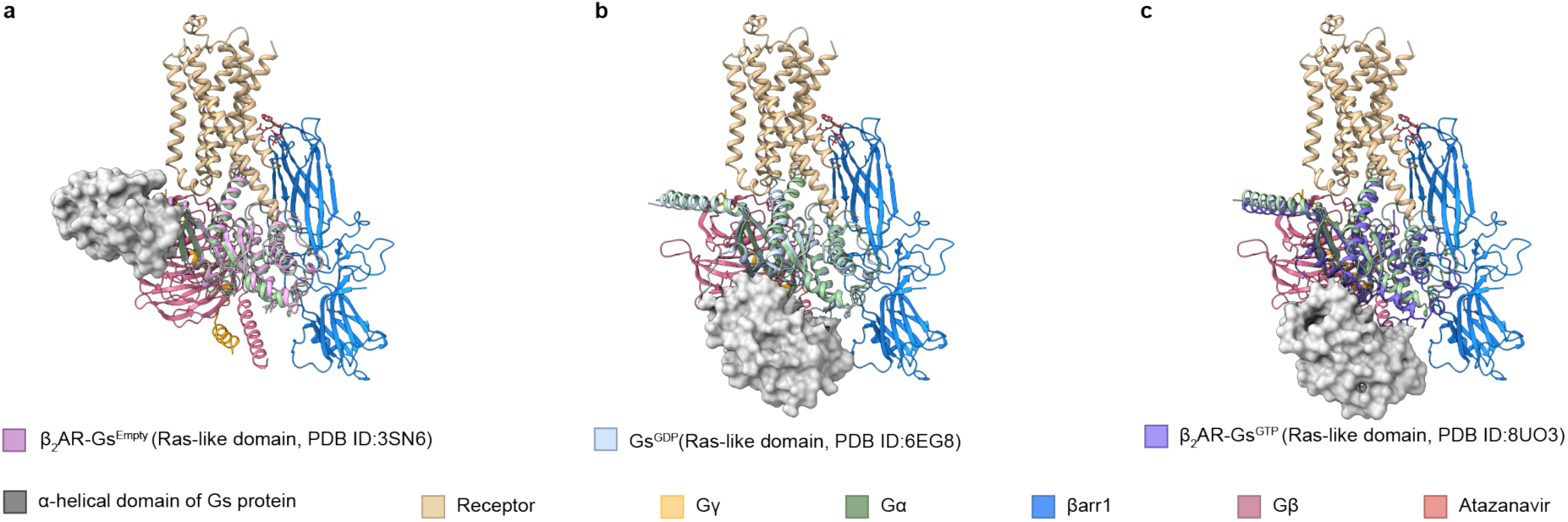
Overlay of megacomplex with different nucleotide-binding states of Gs a-c, Structural alignments of megacomplex with Gs protein in different nucleotide bound states suggesting the binding position of βarr1 (blue) is compatible with each state of Gs protein, including the nucleotide-free state (PDB ID: 3SN6, plum, **a**). the GDP-bound state (PDB ID: 6EG8, light blue, **b**), GTP-bound state (PDB ID: 8UO3, purple, **c**). The α-helical domain (AHD) is shown in gray surface.

**Extended Data Fig. 9.**
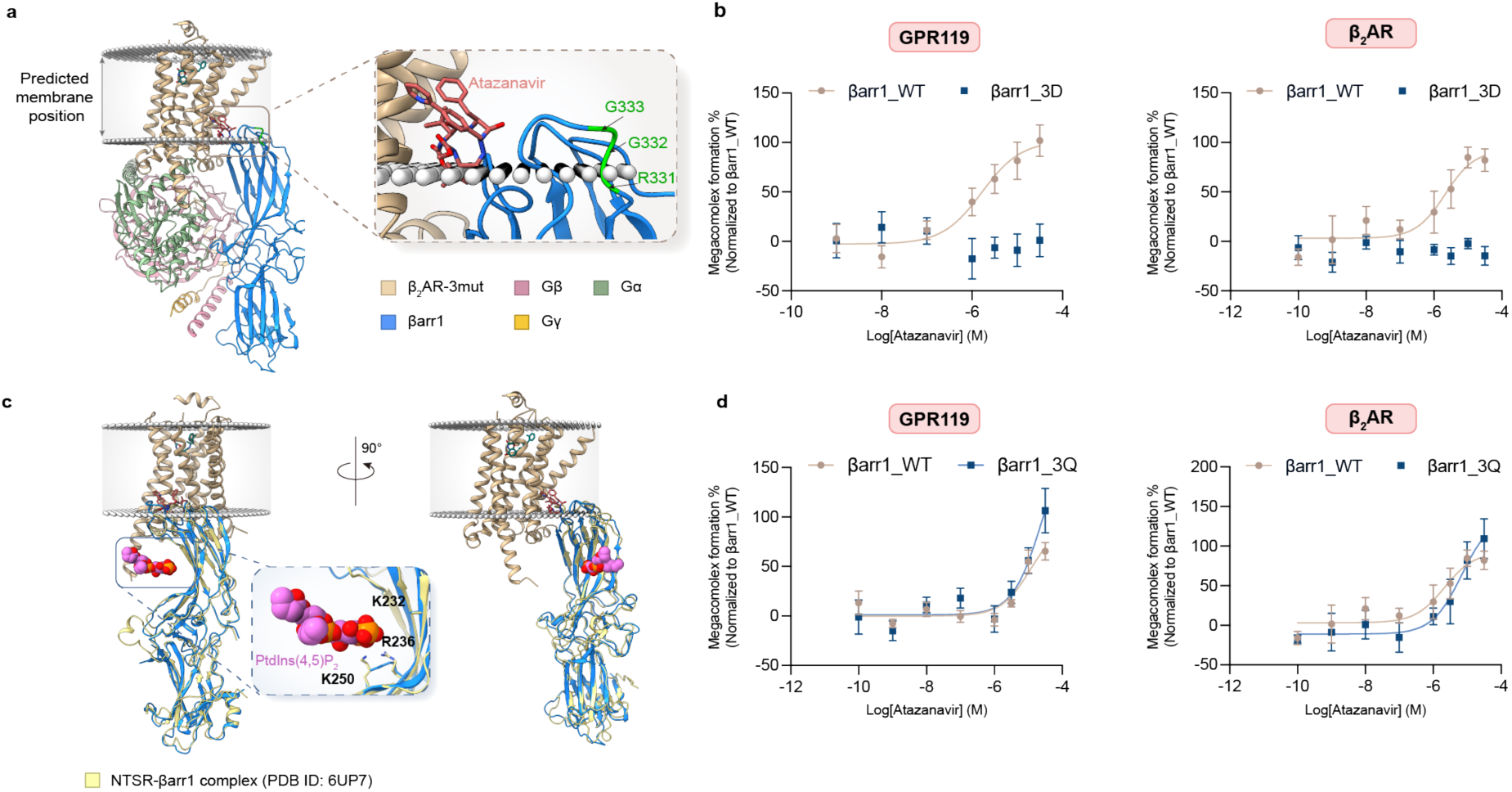
Influence of membrane anchoring and phosphoinositides on the assembly of megacomplex **a,** The C-edge loops of βarr1 penetrate into the membrane bilayer. The position of membrane is predicted by OPM server (https://opm.phar.umich.edu/ppm_server). The three residues (R331, G332 and G333) responsible for βarr1 membrane anchoring are colored in green. **b,** Atazanavir induced megacomplex assembly requires the membrane anchoring of βarr1 for both GPR119 (left) and β_2_AR (right). **c,** Structural alignment suggests the PIP_2_ (magenta sphere) binding site of βarr1(blue) is exposed to solvent in the structure of megacomplex, revealed by superimposition of the NSTR1-βarr1 structure (colored in yellow). **d,** The βarr1_3Q mutant maintains the ability to form megacomplex. Data are shown as Mean ± S.E.M. of 3 independent replicates.

**Extended Data Table. 1.**
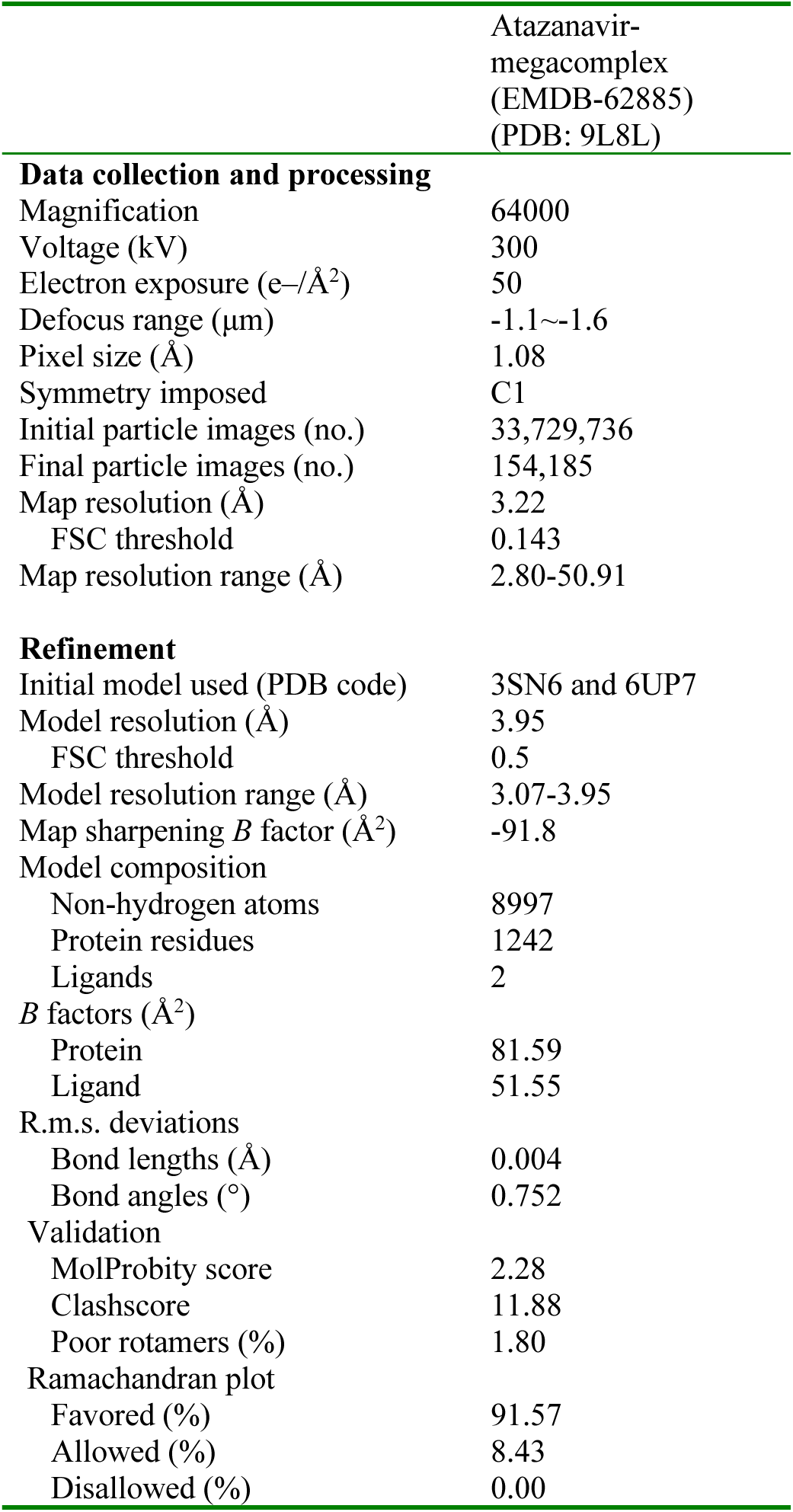
Cryo-EM data collection, refinement and validation statistics.

|  |  |
| --- | --- |
|  | Atazanavir-megacomplex<br>(EMDB-62885)<br>(PDB: 9L8L) |
| **Data collection and processing** |  |
| Magnification | 64000 |
| Voltage (kV) | 300 |
| Electron exposure (e-/Å<sup>2</sup>) | 50 |
| Defocus range (µm) | -1.1~-1.6 |
| Pixel size (Å) | 1.08 |
| Symmetry imposed | C1 |
| Initial particle images (no.) | 33,729,736 |
| Final particle images (no.) | 154,185 |
| Map resolution (Å) | 3.22 |
| FSC threshold | 0.143 |
| Map resolution range (Å) | 2.80-50.91 |
| **Refinement** |  |
| Initial model used (PDB code) | 3SN6 and 6UP7 |
| Model resolution (Å) | 3.95 |
| FSC threshold | 0.5 |
| Model resolution range (Å) | 3.07-3.95 |
| Map sharpening *B* factor (Å<sup>2</sup>) | -91.8 |
| Model composition |  |
| Non-hydrogen atoms | 8997 |
| Protein residues | 1242 |
| Ligands | 2 |
| *B* factors (Å<sup>2</sup>) |  |
| Protein | 81.59 |
| Ligand | 51.55 |
| R.m.s. deviations |  |
| Bond lengths (Å) | 0.004 |
| Bond angles (°) | 0.752 |
| Validation |  |
| MolProbity score | 2.28 |
| Clashscore | 11.88 |
| Poor rotamers (%) | 1.80 |
| Ramachandran plot |  |
| Favored (%) | 91.57 |
| Allowed (%) | 8.43 |
| Disallowed (%) | 0.00 |

